# Complete representation of a tapeworm genome reveals chromosomes capped by centromeres, necessitating a dual role in segregation and protection

**DOI:** 10.1101/2020.04.08.031872

**Authors:** Peter D. Olson, Alan Tracey, Andrew Baillie, Katherine James, Stephen R. Doyle, Sarah K. Buddenborg, Faye H. Rodgers, Nancy Holroyd, Matt Berriman

## Abstract

**Background:** Chromosome-level assemblies are indispensable for accurate gene prediction, synteny assessment and understanding higher-order genome architecture. Reference and draft genomes of key helminth species have been published but little is yet known about the biology of their chromosomes. Here we present the complete genome of the tapeworm *Hymenolepis microstoma,* providing a reference-quality, end-to-end assembly that represents the first fully assembled genome of a spiralian/lophotrochozoan, revealing new insights into chromosome evolution.

**Results:** Long-read sequencing and optical mapping data were added to previous short-read data enabling complete re-assembly into six chromosomes, consistent with karyology. Small genome size (169 Mb) and lack of haploid variation (1 SNP/3.2 Mb) contributed to exceptionally high contiguity with only 85 gaps remaining in regions of low complexity sequence. Resolution of repeat regions reveals novel gene expansions, micro-exon genes, and spliced leader transsplicing, and illuminates the landscape of transposable elements, explaining observed length differences in sister chromatids. Syntenic comparison with other parasitic flatworms shows conserved ancestral linkage groups indicating that the *H. microstoma* karyotype evolved through fusion events. Strikingly, the assembly reveals that the chromosomes terminate in centromeric arrays, indicating that these motifs play a role not only in segregation, but also in protecting the linear integrity and full lengths of chromosomes.

**Conclusions:** Despite strong conservation of canonical telomeres, our results show that they can be substituted by more complex, species-specific sequences, as represented by centromeres. The assembly provides a robust platform for investigations that require complete genome representation.

## Background

Parasitic flatworms are responsible for a significant part of the global worm burden and are ubiquitous parasites of effectively all vertebrate species and many invertebrate groups. Over the past decade reference and draft genomes of key fluke and tapeworm species have been produced including the causative agents of schistosomiasis, neurocysticercosis and hydatid disease [1–6]. Subsequently, improved assemblies and annotations have been published [7] and/or released to the public, as have RNA sequences from an increasing number of transcriptomic studies, profiling genome-wide gene expression for different life cycle stages, cell compartments and experimental conditions [8–11]. Most recently, the diversity of draft genomes of both flatworm and roundworm helminths has been expanded, enabling broader circumscription of helminth-specific gene families and more informative comparative analyses [12]. Despite the growing number of such resources for helminths, little is yet known about their genomic architecture.

Rodent/beetle-hosted *Hymenolepis* species are among the principle tapeworm laboratory models as they enable access to all stages of their complex life cycle. A draft genome of the laboratory strain of the mouse bile-duct tapeworm [13], *Hymenolepis microstoma,* was published in 2013 [6] and updated with additional data and re-released as version 2 on WormBase ParaSite (WBP) [11] in 2015 (details of the v2 assembly are described in [8]). Here we present the third major release of the genome; a reference quality update to the assembly that was made available to the public with the 12^th^ release of WBP (December 2018). The genome has been assembled into full chromosomes, based on the addition of long-read sequence data to previous short-read data followed by extensive alignment, manual review and re-assembly guided by optical mapping data. With this release, *H. microstoma* represents the most completely assembled genome of the lophotrochozoan superphylum.

## Results

### A complete chromosomal representation of the *Hymenolepis microstoma* genome

Using a combination of sequencing technologies we have produced a 169 Mb v3 assembly of the *H. microstoma* genome that is consistent with the known karyotype [14,15]: six scaffolds ranging in size from 17.5 to 43 Mb represent the end-to-end sequences of the six chromosomes (Chr) (Fig. 1, Table S1), while a single, additional contig represents the mitochondrial genome (for a description see Fig. S1). A hybrid assembly was produced based on independent assemblies of long-read Pac-Bio™ sequence data (127x genome coverage), short-read Illumina™ sequence data (115x coverage) and Iris^®^ optical mapping data (77x coverage), and included extensive manual improvements as detailed in the Methods. In total, only 85 scaffolding gaps remain and each is bounded by highly repetitive sequences. Thus collapsed repeats (i.e. tandem repeats assembled as one) rather than novel, non-repetitive sequences likely account for any missing data in gapped regions. The v3 assembly therefore represents an effectively complete picture of the genome both in terms of sequence coverage and assembly and represents a step-change compared with previous releases, with all metrics of assembly contiguity improved by orders of magnitude (Table 1).

**Fig. 1.**
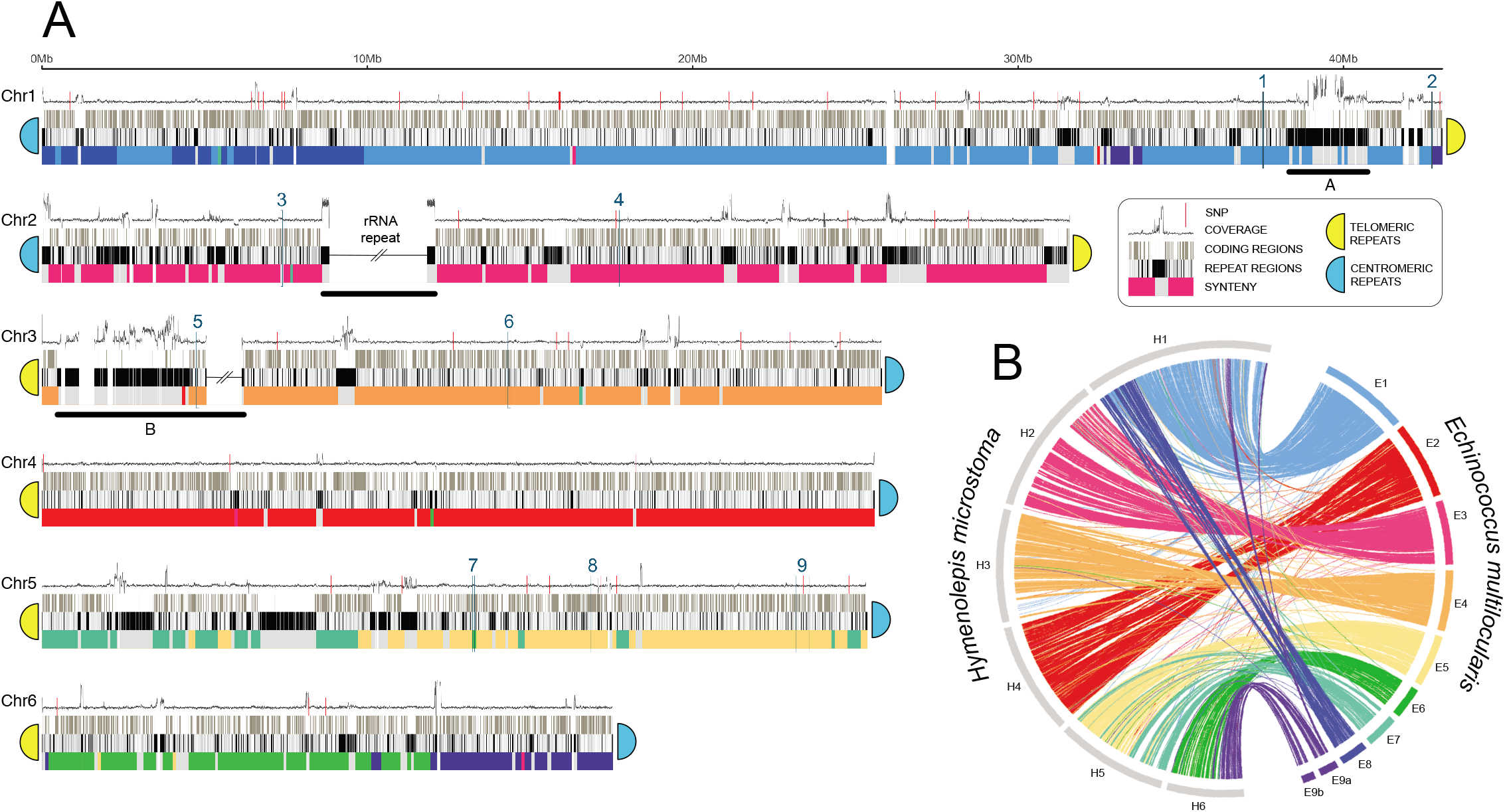
Idiogram of *Hymenolepis microstoma* chromosomes. (**A**) Each chromosome is depicted by three horizontal tracks showing the positions of coding regions, repeats and synteny relative to *Echinococcus multilocularis* (shown in **B**). Synteny is based on 100 kb windows, coloured according to the *E. multilocularis* chromosome with the greatest total number of residues matching using Promer (Methods). Where no hits were found, we coloured the window grey. Above the tracks a graph shows the depth of coverage of Illumina reads mapped against the assembly. Single nucleotide polymorphisms (SNP) shown as red vertical lines along the sequence coverage graph. Red horizontal bars show two interruptions in synteny on Chr1 that reveal a misassembly in the *E. multilocularis* reference genome (see text). Positions of telomeric and centromeric repeat arrays that the chromosome ends are indicated. Regions identified as having enriched pfam clusters are numbered. Regions underscored with horizontal bars and labelled A, B and rRNA depict large repeat arrays discussed in the text. (**B**) shows *H. microstoma* assembly scaffolds aligned against those of *E. multilocularis.*

**Table 1.**
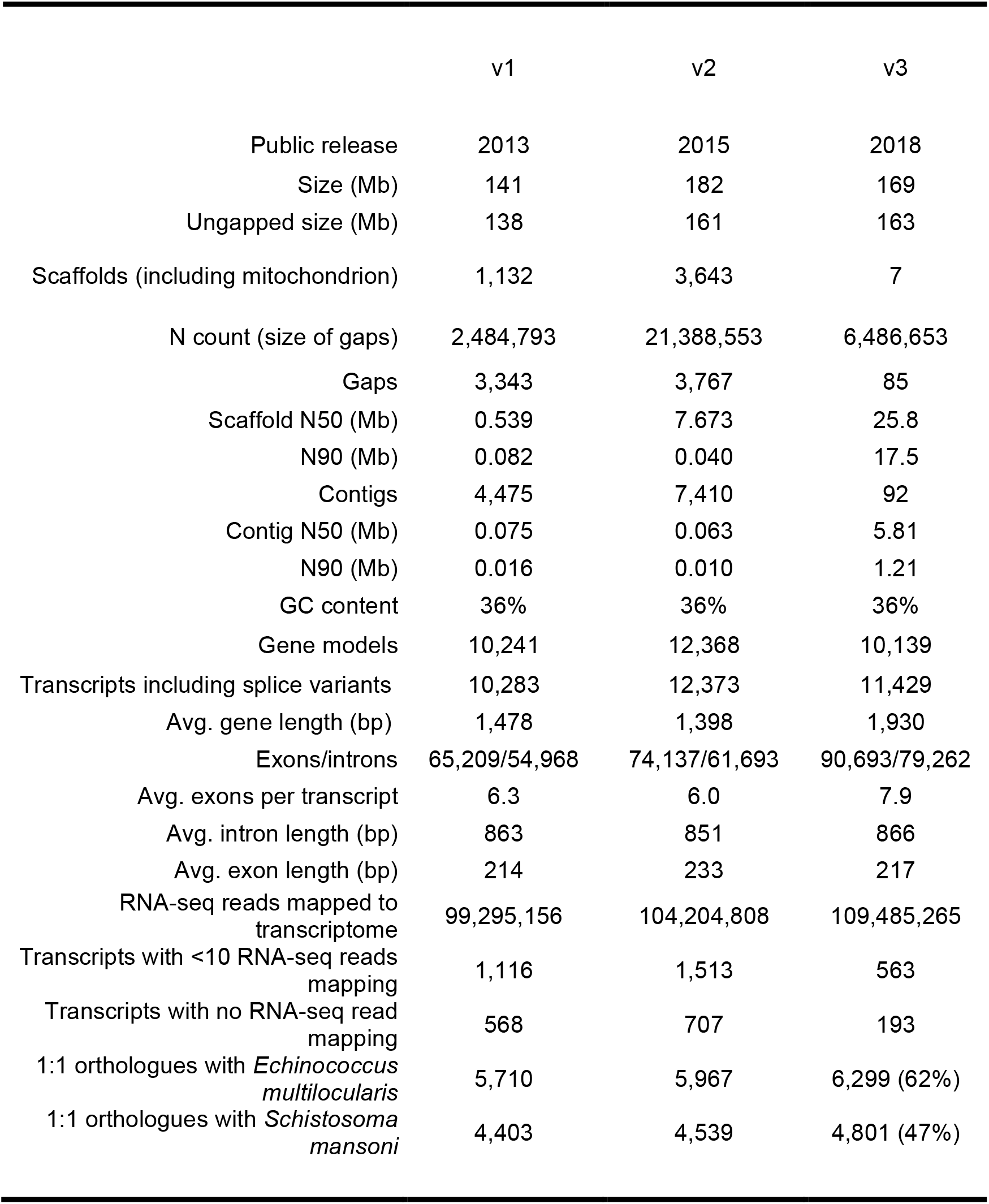
Assembly metrics among *Hymenolepis microstoma* genome releases

|  | v1 | v2 | v3 |
| --- | --- | --- | --- |
| Public release | 2013 | 2015 | 2018 |
| Size (Mb) | 141 | 182 | 169 |
| Ungapped size (Mb) | 138 | 161 | 163 |
| Scaffolds (including mitochondrion) | 1,132 | 3,643 | 7 |
| N count (size of gaps) | 2,484,793 | 21,388,553 | 6,486,653 |
| Gaps | 3,343 | 3,767 | 85 |
| Scaffold N50 (Mb) | 0.539 | 7.673 | 25.8 |
| N90 (Mb) | 0.082 | 0.040 | 17.5 |
| Contigs | 4,475 | 7,410 | 92 |
| Contig N50 (Mb) | 0.075 | 0.063 | 5.81 |
| N90 (Mb) | 0.016 | 0.010 | 1.21 |
| GC content | 36% | 36% | 36% |
| Gene models | 10,241 | 12,368 | 10,139 |
| Transcripts including splice variants | 10,283 | 12,373 | 11,429 |
| Avg. gene length (bp) | 1,478 | 1,398 | 1,930 |
| Exons/introns | 65,209/54,968 | 74,137/61,693 | 90,693/79,262 |
| Avg. exons per transcript | 6.3 | 6.0 | 7.9 |
| Avg. intron length (bp) | 863 | 851 | 866 |
| Avg. exon length (bp) | 214 | 233 | 217 |
| RNA-seq reads mapped to transcriptome | 99,295,156 | 104,204,808 | 109,485,265 |
| Transcripts with <10 RNA-seq reads mapping | 1,116 | 1,513 | 563 |
| Transcripts with no RNA-seq read mapping | 568 | 707 | 193 |
| 1:1 orthologues with *Echinococcus multilocularis* | 5,710 | 5,967 | 6,299 (62%) |
| 1:1 orthologues with *Schistosoma mansoni* | 4,403 | 4,539 | 4,801 (47%) |

### The re-estimated proteome reveals novel gene expansions and previously unidentified classes of genes

The high quality of the genome assembly enabled a more complete complement of genes to be identified. More than 1,700 genes were structurally improved, resulting in an increased average gene length and number of exons per gene despite the total number of models increasing only slightly from the first version (Table 1). In total, 10,139 gene models and 1,310 splice variants were identified using Braker2 [16]. Using Kallisto [17], 10% and 5% more RNA-seq reads map to the v3 transcriptome than to v1 and v2, respectively. Using Orthofinder [18], many transcripts showed clear one-to-one orthology with two near-complete, chromosome-level genome assemblies of other parasitic flatworms: 62% with the hydatid tapeworm *Echinococcus multilocularis* (v4) and 47% with the human blood fluke *Schistosoma mansoni* (v7) (Table 1, Table S2). Compared with the v1 and v2 assemblies, this amounts to 8% and 6% more one-to-one orthologues with *E. multilocularis* and 12% and 6% more with *S. mansoni,* respectively. Overall, the number of genes and average intron and exon size of the v3 proteome is most consistent with the v1 release, whereas the v2 annotation contained an inflated gene count. This indicates that the gene model estimates have stabilized, and together with the assembly and proteome completeness metrics, reflects the advanced level to which the annotation of coding regions has been completed for this genome. A full list of *H. microstoma* gene models and annotations together with *E. multilocularis* orthologues is given in Table S3.

Consistent with the expansion of previously under-represented repeat arrays discussed below, we find that 99 genes previously present as single copies now exist as families with at least three paralogues (Fig. S2, Table S4). Amongst the 12 families with the largest expansions (≥ 5-fold) compared with the v1 genome, a notable example is a C2H2-type zinc finger gene that now has ten copies where previously there was just one. Three families (encompassing 16 genes in v3 but only 3 in v1) are similar to major vault proteins – a cytoplasmic ribonuclear protein complex – and seven families have no obvious sequenced-homologs in other organisms and potentially represent proteins with novel biological functions.

Using the Benchmarking Universal Single-Copy Orthologs (BUSCO) approach [19], 77% of expected genes were identified as complete and without duplication (Table S5). This compares favourably with the manually finished reference genomes of *E. multilocularis* (70%) and *S. mansoni* (73%); completeness scores for parasitic flatworms always fall considerably short of the 100% benchmark. It is therefore likely that many suggested ‘core’ metazoan genes have been lost or have significantly diverged in the flatworm lineage, rather than being erroneously absent from these assemblies. For example, of the 178 BUSCO core genes missing from the v3 assembly, 160 are also missing from *E. multilocularis* and 135 from *S. mansoni* (Table S6). Another factor is likely to be that the lophotrochozoan superphylum is represented by only three species in the BUSCO metazoan database (v3.0.2: two molluscs and one annelid worm). Such under-representation of one of three superphyla may be biasing the circumscription of ‘core’ genes in the Metazoa.

Previously generated RNA-seq data representing different life cycle stages and regions of the adult, strobilar worm were re-mapped to the new v3 assembly and proteome and the resulting table of counts used to estimate differentially expressed genes as described in Olson et al. [8]. Complete lists of up/down-regulated genes ranked by their log2 fold-change are given for all sample contrasts in Tables S7.1-7.7. Comparison with estimates based on the v2 assembly reported in Olson et al. [8] shows a highly linear relationship with the new estimates (Fig. S3) and tight clustering among sample replicates based on principal component analyses (Fig. S4A). Heat map analyses (Fig. S4B) indicate that the transcriptome of the scolex-neck region of the adult is more similar to that of the metamorphosing larvae than to the mid or end reproductive regions of the adult, and this was also shown to be supported by subsets of genes representing signalling pathways and transcription factors as discussed in [8]. Thus while the new analyses supersede those in [8] and include additional differentially expressed genes new to the v3 proteome (highlighted in Tables S7.1-7.7), they also corroborate our previous inferences of differential gene expression.

### Transposable elements comprise a quarter of the genome

Transposable elements (TEs) are among the principal drivers of gene evolution and genome architecture and often comprise the bulk of the DNA in many organisms [20]. TEs comprise approximately 23% of the v3 assembly, although as discussed below the true proportion is likely to be even greater. Of the 23%, 1% is derived from Long Interspersed Nuclear Elements (LINEs), 2% from Long Terminal Repeat retrotransposons and 4% from DNA transposons (Table S8), the most common of which are Mariner-like elements. Although most TEs are highly dispersed, many exist in either a small number of locations or a single location in the genome (Fig. 2). For example, there is a single island of Ginger-type DNA transposons (Chr5: 18.2–18.4 Mb), L1 elements are concentrated on Chr2 (15.4–16.2 Mb) and L2 elements are concentrated on Chr5 (2.2–6.4 Mb). 14.8% of the total repetitive sequence remains unclassified (Fig. 2, Table S9).

**Fig. 2.**
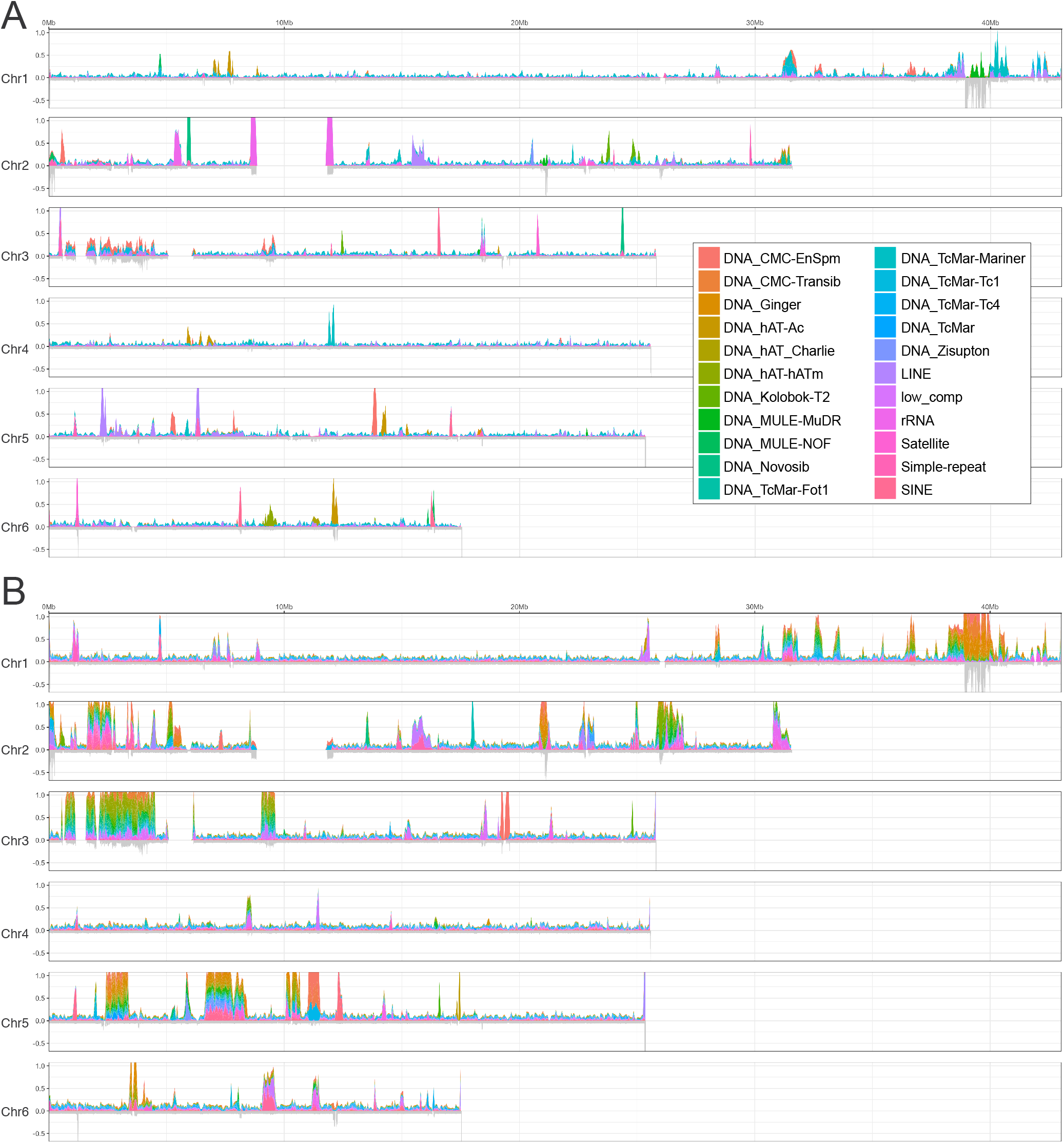
Distribution of transposable elements. (**A**) Transposable elements classified by RepeatModeller (v1.0.11). (**B**) Additional unclassified elements.

Although the addition of long read data in the present assembly enabled full resolution of many more repeat arrays than in previous versions, the depth of coverage of reads realigned to the genome assembly is inordinately high in many places (Fig. 1) indicating that for some repeats, multiple sequenced copies are aligning to fewer copies in the assembled consensus. The true size of some of the largest repeat arrays therefore remains under-represented, including the ribosomal RNA, telomeric and centromeric arrays. Two of the largest examples are on Chr1 (38.9–40.7 Mb) and Chr3 (0.75–4.2 Mb) that are currently assembled into sequences less than half of their expected size based on the relative depth of coverage (labelled A and B, respectively, on Fig. 1). In contrast, Chr4 is notable in having a low proportion of repeats; only 14% of the chromosome is classified as repeat compared with 21-28% across the other chromosomes. The ribosomal RNA array located on Chr2 stands out as the most prominent single repeat type, with an assembled length of 767 kb (0.45% of the assembly). However, its true size based on depth of sequence coverage is likely to be closer to 7.5 Mb (4.4% of the genome), further discussed below.

Repeat content in the first published tapeworm genomes was reported at 7-11%, of which only 2% was attributed to TEs [6]. This proportion of repeats and TEs is exceptionally low and was most likely a reflection of both the inability to fully resolve repetitive regions using shortread data and differences in the identification of TEs. Although TE content is highly variable both across and within animal taxa [21], estimates here of ~25% of the genome content is more typical of metazoans in general and closer to that reported for *S. mansoni* (~35%) [1].

### Variable repeat regions explain length discrepancies in sister chromatids

It was noted from karyology that sister chromatids are not equal in length [14] and that this was especially visible in the largest pair [15]. Although these studies could not rule out the possibility that such differences resulted from the squash technique employed, our sequence data corroborate their observations; whereas we see little to no sequence variation in our assembled contigs, optical mapping data suggest that the largest tandem repeats, which remain elusive to full resolution, could have differing lengths in each pair of sister chromatids. For example, while an optical contig spans the rRNA repeat on Chr2 (the second largest chromosome), giving a short 200 kb form with 17 copies, another optical contig extends into but not across the array, and likely represents the longer version of a larger, alternative haplotype (Fig. S5). It is not possible to directly measure the length of this latter copy but using mapped coverage of Illumina reads from a single library, Chr2 has a median coverage depth of 96x, yet there is a median coverage of 754x over the 486 kb region containing the repeat. We therefore extrapolate that the repeat region exists in the sister chromatid as sequence close to 7.5 Mb. Thus sister chromatids from Chr2 could vary in length by ~25% due to dimorphism in this one repeat region alone. Several other less extreme cases of optical contigs giving two different lengths for the same locus are apparent in the whole genome optical map (Fig. S6), and there are other large repeat regions whose full size is not currently known that could contribute further to homologous chromosomes having unequal lengths.

### Micro-exon genes are identified in the v3 assembly

Genes containing micro-exons that code for as little as a single amino acid occur throughout biology [22]. However, the term micro-exon gene (MEG) was coined for a class of gene that was first identified in the genome of *S. mansoni* [1] and subsequently in *E. multilocularis* [6]. In these genes, multiple micro-exons are present with lengths divisible by three bases, enabling the creation of proteins varying by a single amino acid via exon skipping [23]. Due to their small exons, MEGs are a challenge for gene-finding and RNA-seq reads often fail to align. In contrast to 72 reported MEGs in *S. mansoni* (we now find 109 in the v7 release) and ≥ 8 in *E. multilocularis* (we now find 35 in the v4 release), none was originally reported for *H. microstoma*. However, the greatly improved assembly and proteome enabled us to identify 52 MEGs with a total of 91 transcripts (Table S10). Ten of the MEGs with 14 transcripts are found in a single region of Chr6 (2,643,059-3,072,453) and all share a conserved amino acid sequence motif (consensus: MRLFILLCFAVTLWACPKQCP) that indicates that they belong to a single gene family that expanded via tandem duplication (Fig. S7). A concerted effort to identify and curate MEGs across several flatworm lineages is a high priority for trying to find clues to the functional roles of this numerous yet poorly understood class of genes. However, as many MEGs contain repetitive sequences they are a challenge to analyse without extensive manual curation and at present orthogroups can not be determined with confidence.

### RNA-seq data demonstrate evidence of spliced leader trans-splicing

Spliced-leader (SL) trans-splicing is an mRNA maturation process in which a 5’ donor sequence encoded by its own locus (i.e. the splice leader gene) is spliced to the 5’ exons of other gene transcripts and was first identified in tapeworms by Brehm et al. [24]. We identified the presence of SL trans-spliced transcripts in the transcriptomes of adult and larval *H. microstoma* for the first time. We hypothesised that leader sequences would be present in total RNA-seq libraries and identifiable by their abundance in soft-clipped read segments after alignment to the genome. Using this approach we successfully recovered the previously identified *E. multilocularis* and *S. mansoni* SL sequences [24,25] from analyses of publicly available RNA-seq libraries (Fig. S8A). Our method identified 3,876 genes as being putatively trans-spliced in *S. mansoni* on the basis of having at least one SL-associated read across all of the libraries analysed, reducing this to a conservative set of 1,219 genes with at least ten SL-associated reads. This is comparable with previous estimates of trans-splicing in *S. mansoni* based solely on total RNA-seq libraries [25]. For *E. multilocularis,* 1,609 genes were identified with ≥ 1 SL-associated read and 527 with ≥ 10 reads.

Clustering soft clipped read segments from *H. microstoma* resulted in three abundant clusters, referred to as SL1, SL2 and SL3 (Fig. S8A). Screening these 23-27 bp putative SL sequences against the genome showed that the SL1 motif is found in each of the two exons that comprise gene model HmN_002290900 (Chr1), SL2 is found in an intronic region associated with gene model HmN_000738800 (Chr3), and SL3 is found in a single exon associated with gene model HmN_000738800 (Chr1). No other region in the genome contained these sequences. Based on these SL sequences we identified 1,341 genes with ≥ 1 read and 496 genes with ≥ 10 reads as being putatively trans-spliced. Of the latter, 449 were associated with all three SL sequences, having at least one read of each SL aligned. Similarly, the total number of trans-spliced transcripts found for each SL was highly similar (SL1 = 18,831, SL2 = 18,725, SL3 = 19,241) and the use of ‘interchangeable’ alternative SL forms was also reported for *E. multilocularis* [6]. Using the annotation tool Apollo [26], we validated a subset of these genes as being trans-spliced based on a sharp drop in RNA-seq coverage at the 5’ end of the gene accompanied by an abundance of soft clipped reads, and by the presence of a consensus splice acceptor (‘AG’) coincident with the accumulation of soft clipped reads (example shown in Fig. S8C). A complete list of trans-spliced gene models and associated SLs found in each RNA-seq sample replicate is given in Table S11. Notably, we found that libraries derived from larval *H. microstoma* samples had five times as many trans-spliced genes as libraries derived from adult worms (Fig. S8B).

Early reports of SL trans-splicing in trypanosomes, nematodes and flatworms led to the mechanism being associated with parasitism and interest in it as a potential novel target for chemotherapy [27]. However, further investigation has continued to expand the range of free-living eukaryotic groups in which it is found and this together with structural and functional similarities in the trans-splicing machinery point to it being an ancient process that has been lost independently in most metazoans [28] rather than a process that has been re-invented numerous times [29]. *H. microstoma* genes identified as being trans-spliced (>= 10 aligned reads) were assigned to 494 orthogroups and in 337 of these cases an *S. mansoni* or *E. multilocularis* gene in the same orthogroup was also identified as being trans-spliced, while a core group of 134 orthologues was found to be shared by all three species (Fig. S8D). Spliced leader trans-splicing has also been identified in free-living flatworms [30], but a full inventory of trans-spliced genes in their genomes is needed to investigate to what extent, if any, the process could be associated with parasitism in the phylum. In *H. microstoma* we found that trans-splicing predominates during larval metamorphosis, a period that has been suggested to represent the phylotypic stage of the tapeworm life cycle [31], suggesting that the process may be associated evolutionarily with ontogeny.

### Comparative analysis of chromosomal synteny reveals evidence of ancient linkage groups

Extensive conservation of synteny is clearly evident when comparing the three chromosome-level assemblies of parasitic flatworms. Large regions of *H. microstoma* align to single, often chromosome-sized regions in *E. multilocularis,* enabling the *H. microstoma* chromosomes to be ‘painted’ based on their *E. multilocularis* equivalents (Fig. 1). Between them there are three breaks in overall synteny and when the tapeworm genomes are compared to the blood fluke further breaks in synteny can be discerned that define blocks of chromosomal regions that have persisted as ancestral linkage groups (Fig. 3), recently termed ‘Nigon units’ [32]. Using *S. mansoni* as an outgroup, we can infer that the three tapeworm breaks in synteny are fusions (H1 *cf.* E1+8, H5 *cf.* E5+7, and H6 *cf.* E6+9) as the synteny blocks that have fused to make these *H. microstoma* chromosomes exist separately in the blood fluke (Table S12). In addition to three fusion events, synteny evidence allows us to unambiguously order and orientate two scaffolds from the *E. multilocularis* assembly to form a single chromosome, corresponding to a single ancestral linkage group (labelled E9 in Fig. 1B and G in Fig. 3C). By doing so, the *E. multilocularis* genome assembly resolves to n=9 chromosomes, in agreement with its karyotype [33].

**Fig. 3.**
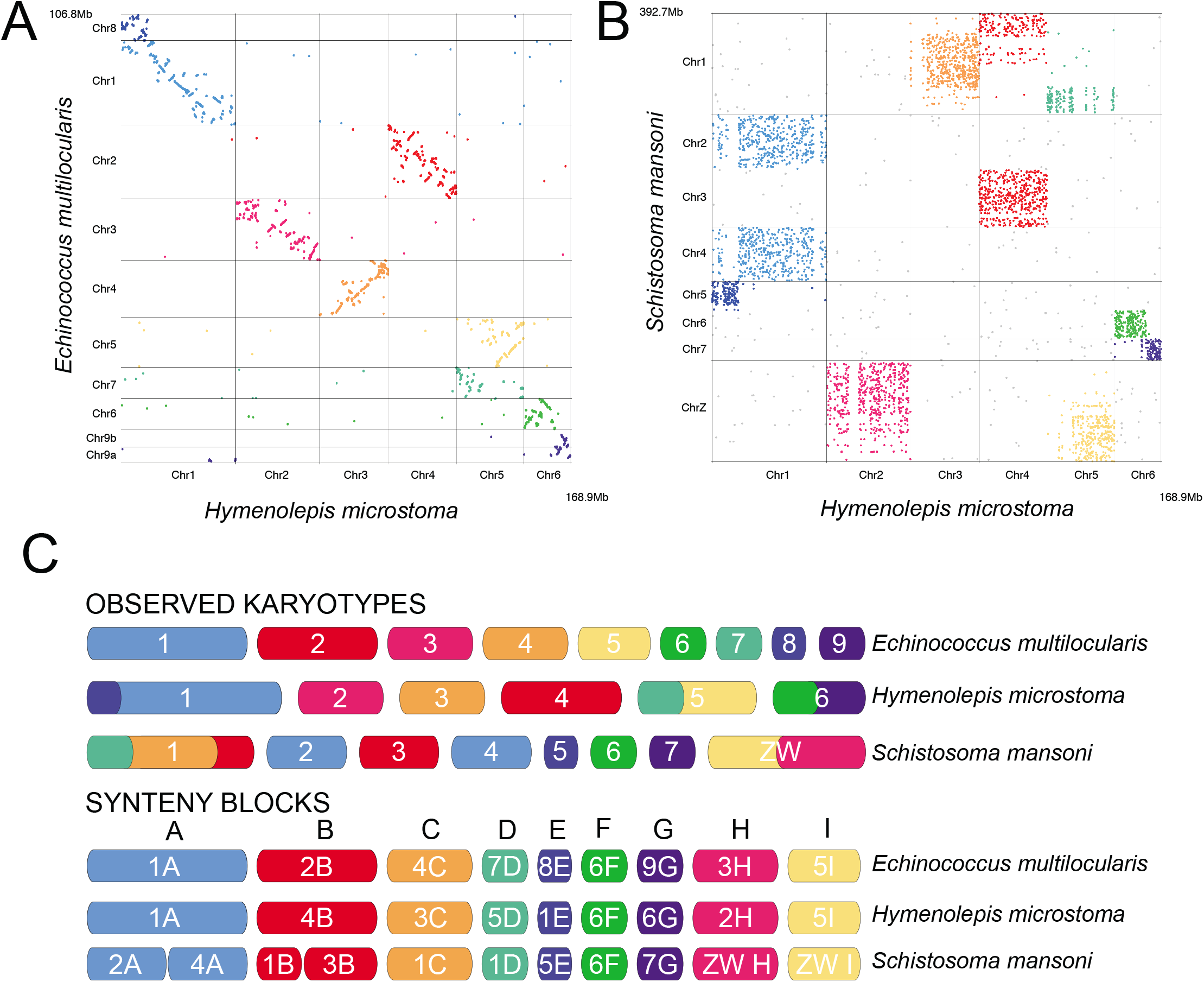
Chromosomal synteny among parasitic flatworms. Comparison between the tapeworms *Hymenolepis microstoma* and *Echinococcus multilocularis* (**A**) shows a high level of synteny not only of scaffold occupancy among the chromosomes, but also of their arrangement within chromosomes, as indicated by their positions arrayed along the diagonal. Comparison between tapeworms and the human blood fluke *Schistosoma mansoni.* (**B**) shows a high level of conservation among chromosomes, but within chromosomes there is little apparent synteny among the scaffolds. In (**C**) their chromosomes are represented by the deduced ancestral linkage groups (‘Nigon’ units) from which we infer that the *H. microstoma* karyotype resulted from the fusion of individual chromosomes still present in *E. multilocularis* and *S. mansoni.*

Although synteny blocks are preserved between these genomes, extensive rearrangements appear to have happened since the fusions occurred which have caused mixing of the synteny blocks such that, in each case, there is no single fusion point, but rather large regions that attest to the fusions. Analysis of one-to-one orthologues reveals that their intrachromosomal order and relative positions are almost entirely scrambled between the blood fluke and tapeworms (Fig. 3B). However, between the two tapeworms we see much greater preservation of gene order, where in some cases (e.g. Chr3 of *H. microstoma* and Chr4 of *E. multilocularis*) effectively no large scale rearrangement has occurred (Fig. 3A). Given that inter-chromosomal rearrangements are exceptionally rare compared with intra-chromosomal rearrangements, the level of shuffling between ancestral blocks provides some indication of the time in which these blocks have been linked together.

### Chromosome ends are capped by a combination of telomeric and centromeric repeats

One of the most striking features of the assembly is that the chromosomes possess telomeric repeats at only one end, whereas opposing ends terminate with a novel repeat array. At the telomeric ends, five of the chromosomes exhibit the canonical hexamer sequence of most telomeres (GGGATT) [34], whereas Chr4 exhibits variation in sequence with the dominant hexamer having a single base variant (TT**C** GGG). At opposing (non-telomeric) ends we find a novel repeat with a median unit length of 179 bp that exhibits several unique traits typical of centromeres: its size is consistent with centromere repeat monomers tending to be about that of one nucleosomal DNA unit (146 bp) [35], *(Homo sapiens,* 171 bp; *Arabidopsis thaliana,* 178 bp; and *Zea mays,* 156 bp.); its sequence is species-specific and highly conserved across chromosomes [36] (with the exception of Chr2 discussed below); and there is only one, large repeat array per chromosome. Moreover, among the sequences that contain this repeat we only find a single junction from unique sequence into the repeat and no junction out of it into another sequence as we find in all other repeats in the genome, and hence it represents a terminal sequence. Finally, we note that in each chromosome the orientation of the repeat remains constant relative to the telomere. That is, by aligning the chromosomes by their telomeric ends (requiring reverse complimenting of Chr1 and Chr2; see Fig. 1) the centromeric sequences are also in alignment. Using the first published assembly [6] and purely algorithmic means (i.e. high copy number, large tandem repeats), this same motif was independently predicted to be the centromere by Melters et al. [37]. We estimate the total size of each repeat array to be at least 5.5 Mb.

Whereas five of the chromosomes have identical motifs, Chr2 contains not only the same novel centromere motif but also a second dominant motif (Fig. S9). In addition, the array is larger and interspersed with other repetitive elements (e.g. gag pol polyprotein) and has a larger sub-telomeric region (Fig. S10). To corroborate our results we used chromosomal fluorescent *in situ* hybridisation (FISH) with probes against the canonical telomeric sequence, showing that only one telomere array is present on each chromosome (Fig. 4A) and that it is opposite to the joined ends of sister chromatids (Fig. 4B), as predicted by our assembly.

**Fig. 4.**
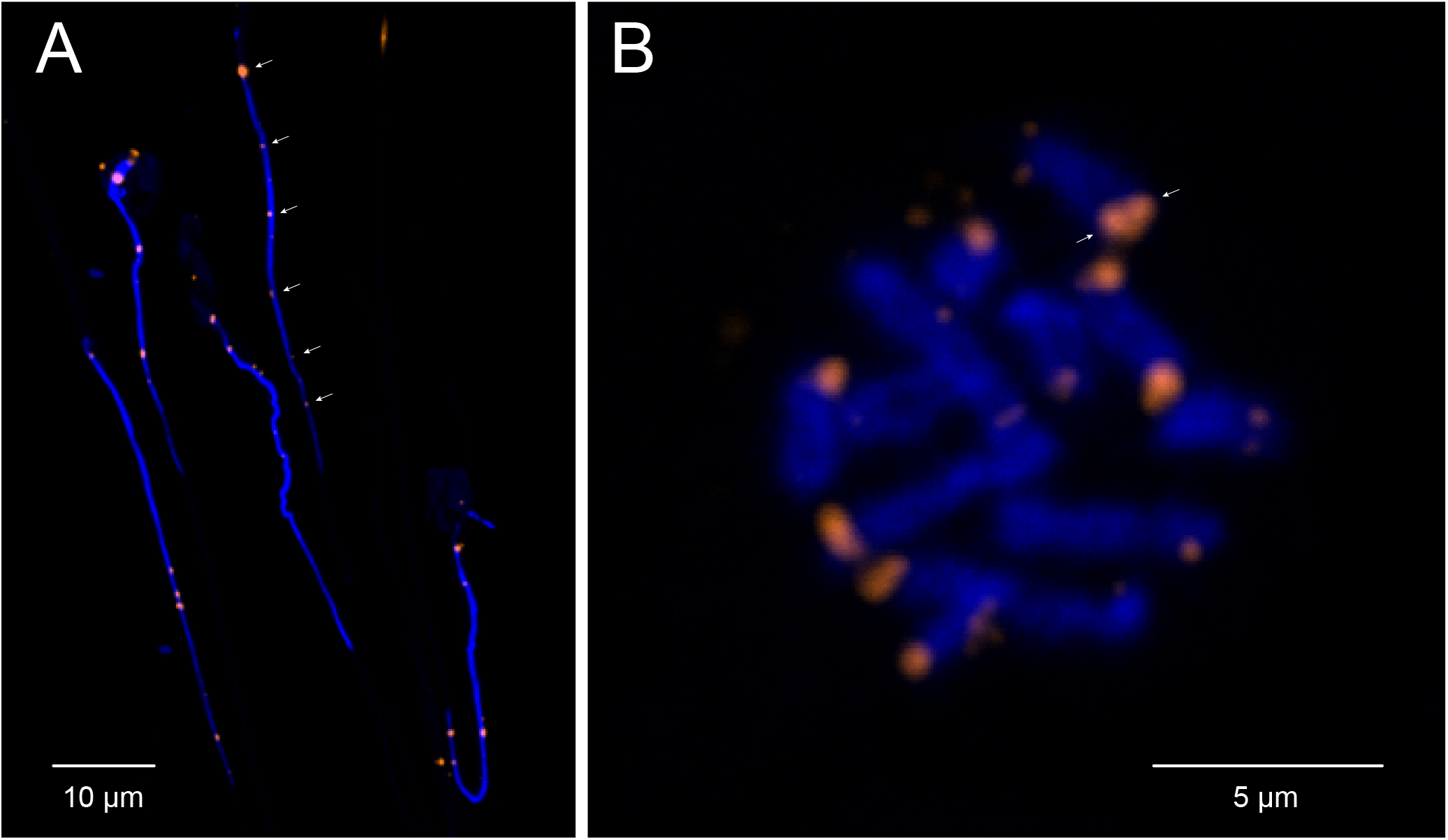
Chromosomal FISH of telomere repeats. Both panels show chromosomal fluorescent in situ hybridisation using probes against the canonical telomere sequence (TTAGGGx7). (**A**) In haploid spermatozoa only one foci is visible for each of the six chromosomes (arrows), whereas two foci per chromosome (= 12) would be expected if telomeric repeats were present on both ends. (**B**) A metaphase figure shows chromatids joined at their centromeric ends, which lack probe signal, whereas probe is visible at the opposing ends of each sister chromatid (arrows).

## Discussion

Such a highly resolved assembly is still unusual and is a product of not only long-read sequence data and optical mapping but also a process of manual improvement. Using Gap5 [38], we were able to scrutinise sequence assemblies from the level of individual base pairs up to whole chromosomes, facilitating diagnosis and resolution of mis-assemblies as well as enabling further scaffolding from clues contained in the read coverage and read-linking data. In this way we have, unusually, been able to place all of the generated sequence data into a chromosomal location, leaving an assembly that is resolved into the same number of scaffolds as the karyotype, with a combined coverage of over 300x. Moreover, although 85 gaps remain there is strong evidence that no novel, complex sequence is missing from the assembly. Assembly was further aided by exceedingly low levels of haploid variation, with only 52 SNPs present in the entire genome. Such low intraspecific genetic variation is very unusual and is presumed to be the result of sequencing a highly inbred laboratory strain [13].

Chromosomes with terminal centromeres have not been demonstrated previously. However, in describing the *H. microstoma* karyotype Hossain and Jones [15] stated that while “the location of the centromere is not clearly visible in the metaphase chromosomes, from the observations of early anaphase of first cleavage it is obvious that all centromeres are terminal or very nearly so.” Here using deep sequencing we demonstrate that the chromosomes do indeed terminate in centromeric arrays that through the course of evolution have most likely come to replace previously existing telomeric arrays. Species lacking canonical telomeres have been found to have chromosomes terminating in either mutated versions of the telomeric sequences themselves (e.g. chironomid midges [39]) or in mosaics of identifiable TEs (e.g. *Drosophila melanogaster* [40]). The 179 bp motif of *H. microstoma* is 30-fold larger than the canonical telomere motif making it unlikely to have evolved directly from a telomeric array. It is also unique, showing no match to known TEs or indeed to any known sequence in the nr database. Thus while definitive validation relies on evidence of centromere-specific histone proteins (CENP-A/CENH3) at the putative region of the chromosome [41], all evidence is consistent with the repeat motif representing the centromere, as independently concluded by Melters et al. [37].

Telomeres are normally present on both ends of chromosomes where they function to maintain linear integrity and length homeostasis [42]. The terminal position of the centromeres suggests that they must act not only as centromeres in providing a substrate for spindle formation during segregation, but that they also play the role of telomeres in protecting chromosome ends from resembling double-stranded breaks. Moreover, being terminal means that the repeats are subject to end replication loss [43] which is normally mediated by a telomerase-dependent replication mechanism [44]. Whether telomeric-specific proteins in *H. microstoma* have evolved to interact with the centromeric motif, or instead a telomeraseindependent mechanism is at play is unknown, but the latter has been suggested as a possibility to explain differences in telomere maintenance between sexual and asexual strains of planarian flatworms [45]. Interestingly, telomere interacting proteins have been found to be under rapid evolution despite strong conservation of their function [42]. This paradoxical observation is similar to the ‘centromere paradox’ in which centromeric sequences are species-specific despite their ultra-conserved role in chromosome segregation [46]. The answer to the paradox appears to be found in the rapid evolution of the sub-telomeric and peri-centrosomal repeats that accompany these arrays [36,42] and it is becoming increasingly clear that despite their functions being perfectly conserved, centromeric and telomeric regions undergo highly dynamic evolution driven by TEs [47].

## Conclusions

Third generation sequencing technologies have enabled the production of highly contiguous genome assemblies that provide more accurate estimates of content as well as the ability to investigate syntenic relationships and other higher-order features of genome architecture. With the third release of the *Hymenolepis microstoma* genome we have produced a reference quality, end-to-end assembly that provides complete chromosomal representation. The hybrid assembly has stabilised estimates of the proteome and non-coding regions and represents a resource effectively free from sampling error. The release thus provides a robust platform to begin systems-level analyses in parasitic flatworms and to this end has been recently used to infer protein-protein interactions based on functional data gathered from major model systems [48].

Producing a fully resolved assembly revealed several unexpected features. Comparative analyses show that large-scale syntenic relationships remain readily apparent even between tapeworms and flukes, which although potential sister groups {Lockyer:2003wj}, represent an ancient split in the Neodermata that was followed by enormous species diversification. Optical mapping indicates that homologous chromosomes differ significantly in length as a result of profound size differences in tandemly repeated arrays of transposable elements and ribosomal genes. Of broadest significance is the finding that chromosomes can terminate in centromeric arrays, providing not only another example of telomere substitution, but also insight into the putative conversion of centromeric motifs. Whether this proves to be a feature unique to this species or is instead common among species with telocentric karyotypes awaits additional chromosome level assemblies of eukaryotic genomes.

## Methods

### Sample preparation

All genome data were derived from the Nottingham laboratory strain [13] of the mouse bile-duct tapeworm *Hymenolepis microstoma* which was maintained *in vivo* using flour beetles (*Tribolium confusum* and *T. castaneum)* and mice. Genomic DNA for long-read sequencing was extracted using a CTAB protocol. 20 mg damp weight of tissue was pooled from the anterior of adult worms (i.e. scolex, neck and immature strobila) which lack reproductive organs or embryos, thereby avoiding genetic variation resulting from gametogenesis and cross-fertilisation. Tissues were homogenised with a plastic pestle in a 1.5 ml Eppendorf, to which was added 0.5 ml CTAB solution (2% w/v hexadecyltrimethyl-ammonium bromide, 100 mM Tris pH 8.0, 20 mM EDTA pH 8.0, 1.4 M sodium chloride, 1% w/v polyvinylpyrrolidone), 50 μl Sarkosyl solution (10% w/v sodium lauroylsarcosinate in 100 mM Tris pH 8.0), 10 μl Proteinase K (20 mg/ml) (ProtK) and 10 μl RNaseA (10 mg/ml). Samples were inverted to mix and incubated at 60°C for 1 hr, after which 0.5 ml Sevac (24:1 chloroform:isoamyl alcohol) was added, the samples mixed and centrifuged at ~13,000 rpm for 3 min. The top, aqueous layer containing DNA was transferred to a new Eppendorf and another 0.5 ml Sevac added and the samples mixed and centrifuged for three minutes. The top layer was transferred to a new Eppendorf, to which 400 μl isopropanol was added and mixed. The samples were centrifuged for 15 min at 4°C, after which the supernatant was removed and 0.5 ml 70% ethanol added. The samples were centrifuged for 5 min at 4°C, the supernatant removed, and the DNA pellet dried in a heating block at 60°C for 5 min. The DNA was re-suspended in 100 μl of ultrapure water and the quantity and quality determined using a NanoDrop spectrophotometer and a TapeStation 2200 fluorometer (Agilent Technologies).

Genomic DNA for optical mapping was extracted from agarose-embedded specimens using the CHEF Genomic Plug DNA kit (BioRad) in order to minimise fragmentation. Four samples were prepared, using 500 and 1,000 larvae (i.e. fully patent cysticercoids harvested from beetles), and 3 (6.6 mg damp weight) and 7 (10.9 mg) sections of adult worm (anterior ~2 cm each; as above). 2% CleanCut (BioRad) agarose was melted at 70°C then cooled to 50°C. Moulds were pre-chilled to 4 °C in the refrigerator. Larval and adult worm sections were left whole and washed in 1 ml phosphate-buffered saline (PBS), then in 200 μl Cell Suspension Buffer, before the latter was added to the washed samples to a final volume of 50 μl. 30 μl of melted agarose was then added and the suspension mixed with a wide bore pipette tip before 80 μl of the agarose-sample mixture was added to a mould well. The mould was then wrapped in parafilm and refrigerated at 4°C for 1 hr. ProtK solution was prepared by adding 16 μl protK stock to 200 μl protK buffer for each 80 μl agarose plug. Refrigerated plugs were removed from their moulds into individual 1.5 ml Eppendorf tubes containing the 216 μl of protK solution and incubated for 2 hr at 50°C in a shaking incubator. The protK was exchanged for fresh solution and the plugs incubated for another 24 hr, after which the protK was exchanged again and the plugs were incubated for another 48 hrs. RNAs were eliminated by treating with 10 μg/ml RNase A (Roche) for 1 hr at 37 °C. Plugs were rinsed briefly three times in Wash Buffer and then four times for 15 min each. ProtK digested specimen plugs were stored in Wash Buffer prior to gDNA recovery.

### Long-read sequencing

19 Gb of long-read sequence data were generated using Pacific Biosciences single-molecule real-time sequencing (SMRT) technology. DNA for sequencing was prepared using the SMRTbell Template Prep Kit 1.0, according to the manufacturer’s protocol, with the exception that shearing was performed using a 26G blunt end needle. A library of ~10 kb sequencing templates was size-selected using SDS-Agarose on a Blue Pippin (Sage Science). Sequencing was performed with the Pacific Biosciences version 2.0 binding kit and sequencing chemistry and a 10 hr runtime, resulting in 1,897,207 raw subreads equivalent to 127x genome coverage.

### Optical mapping

High molecular weight genomic DNA was extracted from *H. microstoma* using the BioRad CHEF Genomic Plug DNA kit as described under sample preparation. An optical map was produced using Bionano Genomics Irys^®^, using the BspQI enzyme. The Irys run generated 40 Gb of data >150 kb that was assembled de novo assembly into 126 contigs with a consensus N50 of 2.4 Mb and coverage of 77x. Hybrid scaffolding of our manually improved Metassembler [49] assembly (below) produced a sequence assembly with 13 scaffolds totalling 165 Mb, along with 7 repetitive scaffolds (4 Mb) that could not be reconciled with the optical map.

### Genome assembly

Two initial *de novo* assemblies were produced using PacBio data: the first used Canu 1.3 [50] and the second used HGAP4 [51], taking the corrected PacBio reads from the Canu assembly process as input. These assemblies were then passed to Metassembler for merging, using the HGAP4 assembly as the primary assembly and the Canu assembly as the secondary assembly. The resulting sequence assembly was passed to Bionano’s Hybrid (optical map) Scaffolder. In addition, an Illumina-only SpAdes assembly was produced [52].

### Manual genome improvement

The genome was manually improved by examining the optical map data in Bionano’s Access software and the sequence data in Gap5 [38]. Errors in the assembly were identified where scaffold breaks needed to be made, or places where new joins could be made. Where groups of Illumina reads mapped to contig ends without their mate-pair, the SpAdes assembly was queried to recover data missing from the assembly. All assembly edits resulting from such investigations were made in Gap5. Soft-clipped reads (PacBio and Illumina) at contig ends were also unclipped where they were found to be in agreement with each other. Many rounds of extending soft-clipped data, remapping, and checking, followed by further extension were undertaken and the results of these incremental improvements were fed back to the Hybrid Scaffolder.

Significant changes to the assembly included breaking an incorrect chromosomal join made by Hybrid Scaffolder and various scaffolding of repetitive scaffolds/contigs. Evidence included repeat junction counting, where repeats were scaffolded, in the absence of reads spanning their entire lengths, if there was only one junction from a non-repetitive region into the repeat at each end. Repeat motifs were analysed with NUCmer [53] and used to determine that many repetitive scaffolds fell into two main repeat types. The two long repeat regions were also joined by analysing their repeat-junctions. Subsequent inspection of these joins (encompassing the last 5 Mb of Chr1 and first 5 Mb of Chr3) in the context of the *E. multilocularis* and *S. mansoni* genomes was used to confirm that they were part of the same chromosome. Most repeat arrays (with the exception of telomeres and centromeres) were located on just one chromosome. A notable exception was a very large repeat occurring as a large complex array on two separate chromosomes; Chr1 around 38-40 Mb and Chr2 around 21-21.2 Mb. Optical contigs failed to bridge either of these repeats and it remains collapsed at both locations. In total there were four junctions from non-repetitive sequence into these repeats. In this instance, a scaffold path was chosen that followed synteny with *E. multilocularis* and *S. mansoni*, given that only three real synteny breaks were found elsewhere.

Extensive optical alignment was used to confirm assembly accuracy (Fig. S6). Apart from three large repeat regions (A, B and rRNA repeat), effectively the entire genome had very good alignment with optical contigs. Some additional gaps remained in the alignments due to large repeats. Optical contigs were much shorter than sequence scaffolds due to a known issue whereby nick sites that occur close together on opposite strands introduce systematic doublestranded breaks that limit the contiguity of Bionano optical maps [54].

This assembly approach yielded the nuclear plus mitochondrial genomes with n = 7 and with 85 sequence gaps remaining, most likely containing repetitive sequence. The mitochondrial contig was circularised to *Cox1* (Fig. S1).

### Gene finding and annotation

Given the fragmented nature of the v1 assembly and questions around the veracity of the v2 annotation set that had 2,000 additional gene models compared with either the v1 gene models or those for *E. multilocularis,* we opted to generate a *de novo* annotation with Braker2 [16] using RNA-seq data as input (for raw data accessions see S1.1 in [8]). RNA-seq reads were mapped to the genome using STAR v2.4.2a [55] and then a merged bam file of these reads was used as input to Braker2. Additionally, Repeat Modeller v1.0.11 [56] and Repeat Masker v1.331 [57] were run and the results used to filter out gene models with >97.5% of their length covered by repeat masked sequence. Annotation was loaded into Apollo [26] and manually assessed. Particular attention was paid to regions of the genome with the highest densities of gene models and it was noted that many of these models had fallen near to, but just below, the 97.5% threshold mentioned above, and upon inspection were generally found to result from incorrect annotation of gene models in tandem repeats and so were removed. OrthoMCL [58] was used to find one-to-one gene mappings between the resulting annotation and the previous v1 and v2 gene models. Where unambiguous mappings were found, the historical gene IDs were transferred and are thus consistent with previous releases. Where mappings were ambiguous or non-existent, new gene IDs were created prefixed with ‘003’ (e.g. HmN_003NNNNNN). The mitochondrial genome was annotated independently using Mitos2 [59].

The distribution of repeats were subsequently analysed using RepeatModeller (v1.0.11) followed by RepeatMasker (v4.0.7).

### Analysis of synteny conservation between flatworms

The *S. mansoni* genome assembly v7 (PRJEA36577) and the latest *E. multilocularis* assembly were obtained from WBP (release 12). Translated alignments of 100 kb windows from each *H. microstoma* chromosome were compared against *E. multilocularis* using Promer v3.07 (--mum setting). Dot plots of synteny based on the position of orthologues was used to further characterise and more accurately determine the position of conserved synteny blocks. One-to-one orthologues were identified between *H. microstoma* and *E. multilocularis* as well as *H. microstoma* and *S. mansoni* using OrthoMCL v1.4 [58]. Each orthologue pair was plotted as a single point and coloured by the genomic location of the *E. multilocularis* and *S. mansoni* genes, respectively.

### Centromere quantification

An attempt was made to quantify the centromeric repeat using Illumina data. One representative unit of the putative centromere sequence (179 bp) and another more specific to the repeat variant found on Chr2 (190 bp) were concatenated with the first 180 bp taken from 50 gene sequences. Using BEDTools [60] coverage, we calculated mean coverage over 10 bp windows for each gene sequence. The median of these mean values taken from all 50 genes was 50.25x. The 179 bp unit had 1,549,563x coverage and the 190 bp unit had 6,237x coverage. From this, we calculated a grand total of 5.5 Mb which we take to be a minimum size estimate for this repeat, in line with the expectation that the centromere repeat is likely to be the largest repeat in the genome [37].

### Variant calling

Variants were called using GATK Unified Genotyper v3.3.0 [61]. The raw variant set was initially filtered to flag variants as low quality if they met the following conditions: quality by depth (QD) < 2; Fisher’s test of strand bias (FS) > 60; RMS mapping quality (MQ) < 40; rank sum of alt versus reference mapping quality (MQRankSum) < −12.5; read position rank sum (ReadPosRankSum) < 8; read depth (DP) < 10. Variants were filtered further using vcftools (v0.1.14) [62] to exclude sites with low quality flags, minimize loci with missing data (“maxmissing 0.8”), exclude indels (“remove-indels”), exclude SNPs with genotype quality (GQ) < 30, and ensure sites were biallelic (“min-alleles 2, max-alleles 2”). Remaining variants were manually curated in Gap5 [38] and a total of 52 were found to be genuine heterozygous calls, giving a SNP rate of 1 per 3.25 Mb. It was subsequently found that these SNPs could be isolated using the following GATK filtering parameters: qual > 120, DP < −4, dels > 55, HaploScore > 45, MapQualRankSum < 1.5, QD > 0.9, SOR > 6, ReadPosRankSum < −2.

### Identification of micro-exon genes

Custom shell and Perl scripts were used to download and parse GFF-formatted annotation from WBP (July 2019) to create a table of exon lengths for each gene. The resulting table was further parsed to identify exons shorter than 70 nucleotides and divisible by three as micro-exons. Genes comprising at least seven exons, with micro-exons constituting at least half of all exons and runs of at least four consecutive micro-exons were deemed to be micro-exon genes. For more information see https://github.com/wbazant/microexons/blob/master/README.md.

### Identification of splice leader sequences and trans-spliced genes

Publicly available RNA-seq libraries were used to identify splice leader sequences in *E. multilocularis* (run accessions: ERR337946, ERR337958, ERR337939, ERR337951, ERR337963, ERR337962), *S. mansoni* (run accessions: ERR022872, ERR022877, ERR022878, ERR022880, ERR022881, ERR022882, ERR1674583, ERR1674584, ERR1674585, ERR1674590, ERR1674591, ERR1674592, ERR506076, ERR506082, ERR506083, ERR506084, ERR506084, ERR506088, ERR506090) and *H. microstoma* (ERR225719-ERR225730, ERR337928, ERR337940, ERR337952, ERR337964, ERR334976).

TruSeq3 Illumina adapter sequences were trimmed from RNA-seq reads using Trimmomatic (v0.39) and reads aligned to the genome using STAR (v2.7.3a) with the following parameters: outFilterMultimapNmax 20, alignSJoverhangMin 8, alignSJDBoverhangMin 1, outFilterMismatchNmax 999, outFilterMismatchNoverReadLmax 0.04, alignIntronMin 20, alignIntronMax 1000000, and alignMatesGapMax 1000000. Annotations downloaded from WBP release 14 were provided to guide alignment. Unique alignments were parsed using a custom python script to identify reads that (a) aligned to annotated genes, or within 500 bp upstream, and (b) were soft clipped by more than 5 bp at the 5’ end relative to the annotated gene. These soft clipped sequences from all libraries were then clustered (cd-hit-est v4.7) and three (*H. microstoma)* or one (*E. multilocularis, S. mansoni)* prominent clusters identified as putative splice leader (SL) sequences. Genes associated with clipped SL reads were considered to be putatively trans-spliced. Genomic splice leader loci were identified by aligning SL sequences against the genome using BLAST. Code is available at https://github.com/fayerodgers/trans_splicing.

### Chromosomal FISH

The asymmetric presence of telomeric repeats on the ends of the chromosomes was investigated empirically via chromosomal fluorescent *in situ* hybridisation (FISH). Chromosome spreads were performed based on the methods of Orosová and Špakulová [63]. Adult worms were freshly harvested from the bile-ducts of mice into plastic petri dishes, rinsed in mammalian saline (0.85% w/v NaCl) and incubated in supplemented media with colchicine (Sigma Aldrich) (M199, 20% foetal bovine serum (FBS), 1% sodium choleate, 0.25% colchicine) for 4 hr at 37°C in a 5% CO_2_ atmosphere. They were transferred to distilled water, cut into pieces, pierced, and incubated for 20 min to allow the cells to swell. The swollen tissues were fixed in Carnoy’s fixative (3:1 methanol:acetic acid) for 30 min and then stored in fixative at 4°C until used 24-48 hr later. A small piece of worm (~1 mm) was put on a microscope slide and 15 μl cold acetic acid added before macerating the piece with needles. Slides were placed on a 45°C hotplate and the cell suspension spread with a metal hook. Excess acetic acid was removed by blotting and the slides dehydrated in an ethanol series (70%, 80%, 90% and 100%) before air drying.

The protocol of Guo et al. [64] for chromosomal FISH was combined with tyramide signal amplification (TSA) for increased detection [65]. A 42 bp oligonucleotide based on the canonical telomere repeat ([TTAGGG]x7) was synthesised commercially and then labelled with digoxigenin-11-2’-deoxyuridine-5’-triphosphate (DIG-11-dUTP) using terminal transferase (Roche) according to manufacturer’s instructions. DIG-labelled probe was purified by sodium acetate and ethanol precipitation and re-suspended in 20 μl water. For each slide, 1 μl of probe was mixed with 250 μl hybridisation buffer (50% formamide, 5x saline-sodium citrate buffer (SSC), 100 μg/ml heparin, 1x Denhardt’s solution, 0.1% Tween 20, 0.1% CHAPS, 10 mM EDTA, 0.5 mg/ml bovine serum albumin (FBS), 5% dextran sulphate).

FISH assays were performed both by hand and using an Intavis InsituPro VSi *in situ* robot using 250 μl volumes for each step except probe hybridisation, which used 200 μl. Slides were incubated in hybridisation buffer for 10 min at RT, then 10 min at 70°C. Probe was hybridised at 70°C for 10 min, then cooled to RT and incubated for 12 hr. Slides were washed 6 times for 5 min each with 2x SSC, 0.5x SSC, then TNT (100 mM Tris-HCl, 150 mM NaCl, 0.1% Tween20). They were then incubated with TNB (5% FBS in TNT) for 15 min before incubation with peroxidase-conjugated anti-DIG antibody (DIG-POD, Roche) 1:200 in TNB for 2 hr at RT. Slides were washed 6 times for 5 min with TNT, then twice each in phosphate buffered saline (PBS) with 0.1% Tween 20 and PBS with 0.1 M imidazole. Signal detection was performed by incubating in rhodamine-conjugated TSA mix (988 μl PBS with 0.1 M imidazole, 10 μl 0.1% H_2_O_2_, 2 μl rhodamine-conjugated tyramide) for 5 min, then washed 6 times for 5 min each in PBST then TNT. Slides were lastly incubated in 1 μg/ml DAPI for 15 min before being washed twice with TNT. The full InsituPro method is given in Additional file 2. Slides were removed from the robot and mounted with coverslips in 87.5% glycerol, 2.5% DABCO, 10% PBS and 1 μg/ml DAPI. Results were visualised and imaged with a Nikon A1 confocal microscope using a 63x oil objective and Nikon NIS software v4, or a Leica DM5000B epifluorescent microscope using a 100x oil objective and Leica LAS software v4. Images were processed to adjust overall levels using Fiji/ImageJ v2 [66].

## Supporting information

Add. File 1: Supplemental Tables

Add. File 2: Robot method

Add. File 3: Supp. Figures S1-10

## Additional Files

### Additional file 1

Supplementary tables. **Table S1.** Chromosome summary. **Table S2.** Comparison of one-to-one orthologues between assemblies and other flatworms. **Table S3.** Gene model annotations and *Echinococcus multilocularis* orthologues. **Table S4.** Paralogous expansions within orthologue groups predicted using successive *H. microstoma* genome assembly versions. **Table S5.** Assessment of genome completeness based on presence/absence of conserved eukaryotic genes. **Table S6.** Presence and absence of BUSCO orthologues (v. 3.0.2) missing in ≥ one flatworm. **Table S7.1.** Differentially expressed gene models in Larvae vs. Whole Adult RNA-seq samples ranked by log2-fold change. **Table S7.2.** Differentially expressed gene models in Scolex-Neck vs. Mid RNA-seq samples ranked by log2fold change. **Table S7.3.** Differentially expressed gene models in Scolex-Neck vs. End RNA-seq samples ranked by log2-fold change. **Table S7.4.** Differentially expressed gene models in Mid vs. End RNA-seq samples ranked by log2-fold change. **Table S7.5.** Intersect of gene models up-regulated in the Scolex-Neck cf. Mid and End. **Table S7.6.** Intersect of gene models up-regulated in the Mid cf. Scolex-Neck and End. **Table S7.7.** Intersect of gene models up-regulated in the End cf. Mid and Scolex-Neck. **Table S8.** Repetitive elements summary. **Table S9.** Repetitive element hotspots. **Table S10.** Micro-exon genes. **Table S11.** Trans-spliced genes. **Table S12.** Chromosome fusions between *H. microstoma* and *E. multilocularis.* (XLSX workbook; 4.7 MB)

### Additional file 2

Method programme for automated chromosomal FISH using the Intavis InsituPro VSi robot. (DOC; 45 KB)

### Additional file 3. Figure S1

Mitochondrial genome. The 13,919 bp *Hymenolepis microstoma* mitochondrial genome was re-assembled from both short and long-read data, yielding over 1000x coverage. The new assembly resolved the full length of a region involving a tandemly repeated 32 bp motif (cf. GenBank accession AP017665.1). This region is identified as one of three origins of replication-heavy strand (OH-a) by MITOS [59] and an adjacent hairpin-loop region as the origin of replication-light strand (OL). Gene order of ribosomal and protein-coding genes is consistent with the hypothesized ground-plan for the mitogenomes of parasitic flatworms as is the absence of the *atp8* gene [67]. (PDF; 1.2 MB)

### Additional File 4: Figure S2

Repeat hotspots. Chromosomal positions of paralogous gene arrays. Abbreviations: ABCB: ATP binding cassette subfamily B; Akr1b4: Aldo keto reductase family 1 member B4; AP: Alkaline phosphatase; AQP: Aquaporin 4; CREBBP: CREB binding protein; DYNLL: Dynein light chain; EiF2c: Eukaryotic translation initiation factor 2c; ENPP: Ectonucleotide pyrophosphatase:phosphodiesterase; EP45: Estrogen regulated protein EP45; GST: Glutathione S transferase; H3: Histone H3; HSP: heat shock protein; hypo: hypothetical protein; MVP: Major vault protein; PARP: Poly [ADP-ribose] polymerase; PiT: Phosphate transporter; PNP: Purine nucleoside phosphorylase; PP2A: Serine:threonine protein phosphatase 2A; PURA: PUR alpha protein; USP: Universal stress protein; RAD51: DNA repair protein RAD51 homolog; SLC22: Solute carrier family 33; TSP: Tetraspanin; TXN: Thioredoxin; ZNF: zinc finger protein. (PDF; 926 KB)

### Additional File 5. Figure S3

Comparison of differentially expressed genes estimated from RNA-seq counts aligned to the v2 and v3 assemblies and gene models. Plots of log2-fold change show highly linear relationships across all sample comparisons, corroborating previous findings [8]. Only 11 genes (yellow), all with small fold-change values, were found to reverse directionality between assembly versions. (PDF; 6.9 MB)

### Additional File 6. Figure S4

Comparison of RNA-seq sample counts against the v2 and v3 assemblies and gene models. Principle component analyses (**A**) show tight clustering of sample replicates based on counts using both assemblies, while in the v3 (right) the Larvae, Scolex-Neck and Whole Adult samples are arrayed only along PC1, with the transcriptome of the Scolex-Neck mid-way between those of the Larvae and Whole Adult samples. The Mid and End samples are further differentiated from the other samples along PC2. Heatmap clustering (**B**) shows that the transcriptome of the Scolex-Neck region is more similar to that of midmetamorphose larvae than to middle or end regions of the adult worm, as discussed in [8]. (PDF; 267 KB)

### Additional File 7. Figure S5

Optical map contigs aligned to the genome assembly of the rRNA repeat array. Five contigs from the optical map are shown with the segment that aligns to the sequenced repeat indicated by coloured bars. The largest map contig (arrow) represents one haplotype containing the rRNA tandem repeat (pink bar) as well as the left (blue bar) and right (yellow bar) flanking regions. Other optical map contigs either contain the repeat together with either 5’ or 3’ flanking region, and likely represent an alternative haplotype, or have an insufficient amount of unique sequence to unambiguously determine their position within the repeat array. (PDF; 1.9 MB)

### Additional File 8. Figure S6

Whole genome optical maps aligned to v3 assembly. Circled regions show where optical map data indicate alternative haplotypic versions. Regions labelled A and B, together with the rRNA array, represent the largest repeat regions where haplotype differences could account for visible length differences in sister chromatids (see text). Chromosomes are numbered and the positions of the telomeric repeats indicated by red dots. (PDF; 3 MB)

### Additional File 9. Figure S7

Alignment of the N-terminal regions encoded by a tandem array of micro-exons genes located on Chr 6. The shared amino acid motif (consensus MRLFILLCFAVTLWAC) indicates that this gene array evolved through tandem duplication. (PDF; 1 MB)

### Additional File 10. Figure S8

Spliced leader trans-splicing. (**A**) Clustering of sequences soft clipped from aligned RNA-seq reads. The most abundant clusters represent known (*E. multilocularis, S. mansoni)* or candidate (*H. microstoma)* splice leader (SL) sequences which are given in the table below. (**B**) The prevalence of trans-splicing in different life stages and regions of the adult worm. Genes were considered trans-spliced if > 10 SL reads (SL1, SL2 or SL3) aligned across all libraries analysed. Of these genes, plot represents instances of at least one SL read aligning in each sample. Note that there are 5x as many genes trans-spliced in larvae than in the adult samples. Three replicates per sample. (**C**) An example of a gene (HmN_000032200) that is trans-spliced in larval but not adult samples, visualised using Apollo. Left: track 1 shows a coverage plot of all aligned reads; track 2 represents alignments of uniquely-mapping soft-clipped reads (soft clipping represented by a thick blue bar at the end of the read). Arrow indicates accumulation of soft clipped reads at proposed SL-acceptor site. Right: Coverage plots of all aligned reads in three larval and three adult libraries. Arrows indicate proposed SL-acceptor sites present in the larval but not adult libraries. (**D**) Venn diagram of trans-spliced orthogroups shared between parasitic flatworms. (PDF; 5.6 MB)

### Additional File 11. Figure S9

Multiple alignment of the terminal centromeric repeats of each chromosome. 26 consecutive repeat copies were taken from a single location at the end of each of the six chromosomes in turn and aligned in order (top 26 = Chr1, next 26 = Chr2 etc.). Strong conservation of the 179mer centromeric repeat is seen across all chromosomes except Chr2 which shows a second novel repeat type. However, searching within the whole of the Chr2 repeat array shows that the ‘canonical’ 179mer observed in the other aligned reads is found with 100% coverage and identity. The terminal array on Chr2 is also much larger than those of the other chromosomes and is interspersed with various other repeats not shown here. Full assembly of the Chr2 terminal array is not resolvable without longer sequencing reads. Notably, when the centrosomal repeat arrays are oriented at the same end of each chromosome their sequences are found to be in alignment. (PDF; 1.7 MB)

### Additional File 12. Figure S10

The terminal centromeric repeat of chromosome 2. A dotter plot shows that the centromeric repeat not only contains a second dominant repeat motif but is also interspersed with other repetitive elements, unlike the other chromosomes that exhibit a tandem array comprised entirely of the novel 179mer. Within the interstitial sequences we find the top blastx hit to Gag-Pol polyprotein, indicating the centromere has been invaded by transposable elements. (PDF; 3.9 MB)

## Declarations

### Ethics approval and consent to participate

Animals were used in accordance with project license PPL70/8684 issued by the UK Home Office to PDO.

### Consent for publication

Not applicable

### Availability of data and material

The datasets generated and/or analysed during the current study are available in the European Nucleotide Archive (www.ebi.ac.uk/ena) under the following accessions; genome assembly GCA_000469805.3, long read sequence data study accession PRJEB2107.

### Competing interests

The authors declare that they have no competing interest.

### Funding

This work was supported by Wellcome (grant 206194) to AT, NEH and MB; BBSRC grant BB/M003949/1 to SRD and BBSRC grant MR/L001020/1 to FHR.

### Authors’ contributions

AT, PDO and MB conceived and designed the study. AT assembled and manually curated the genome and led bioinformatic analyses; AB prepared samples and performed chromosomal in situ hybridisation; KJ conducted differential expression analyses; FHR analysed spliced leader trans-splicing; SRD performed preliminary analyses of synteny and advised on annotation and analytical approaches; NEH coordinated the specimens and sequencing; AT, AB, PDO and MB interpreted results and prepared the paper which was led by PDO and AT. All authors read and approved the final manuscript.

## References

1. Berriman M, Wilson RA, Dillon GP, Cerqueira GC, Ashton PD, Aslett MA, et al. The genome of the blood fluke Schistosoma mansoni. Nature. 2009;460:352–8.

2. Young ND, Jex AR, Li B, Liu S, Yang L, Xiong Z, et al. Whole-genome sequence of *Schistosoma haematobium*. Nat Genet. 2012;44:221–5.

3. Wang X, Chen W, Huang Y, Sun J, Men J, Liu H, et al. The draft genome of the carcinogenic human liver fluke *Clonorchis sinensis*. Genome Biol. 2011;12:R107.

4. Young ND, Nagarajan N, Lin SJ, Korhonen PK, Jex AR, Hall RS, et al. The *Opisthorchis viverríni* genome provides insights into life in the bile duct. Nature Communications. Nature Publishing Group; 2014;5:1–11.

5. Olson PD, Zarowiecki M, Kiss F, Brehm K. Cestode genomics - progress and prospects for advancing basic and applied aspects of flatworm biology. Parasite Immunol. 2012;34:130–50.

6. Tsai IJ, Zarowiecki M, Holroyd N, Brooks KL, Tracey A, Bobes RJ, et al. The genomes of four tapeworm species reveal adaptations to parasitism. Nature. 2013;496:57–63.

7. Protasio AV, Tsai IJ, Babbage A, Nichol S, Hunt M, Aslett MA, et al. A systematically improved high quality genome and transcriptome of the human blood fluke *Schistosoma mansoni*. PLoS Negl Trop Dis. 2012;6:e1455–13.

8. Olson PD, Zarowiecki M, James K, Baillie A, Bartl G, Burchell P, et al. Genome-wide transcriptome profiling and spatial expression analyses identify signals and switches of development in tapeworms. EvoDevo. BioMed Central; 2018;9:1–29.

9. Jex AR, Gasser RB, Schwarz EM. Transcriptomic resources for parasitic nematodes of veterinary importance. Trends Parasitol. Elsevier Ltd; 2019;35:72–84.

10. Grote A, Lustigman S, Ghedin E. Lessons from the genomes and transcriptomes of filarial nematodes. Mol Biochem Parasitol. 2017;215:23–9.

11. Howe KL, Bolt BJ, Shafie M, Kersey P, Berriman M. WormBase ParaSite - a comprehensive resource for helminth genomics. Mol Biochem Parasitol. Elsevier B.V; 2017;215:2–10.

12. International Helminth Genomes Consortium, Coghlan A, Mitreva M, Berriman M. Comparative genomics of the major parasitic worms. Nat Genet. 2018;:1–35.

13. Cunningham LJ, Olson PD. Description of *Hymenolepis microstoma* (Nottingham strain): a classical tapeworm model for research in the genomic era. Parasites & Vectors. 2010;3:123.

14. Proffitt MR, Jones AW. Chromosome analysis of *Hymenolepis microstoma*. Exp Parasitol. 1969;25:72–84.

15. Hossain M, Jones A. The chromosomes of *Hymenolepis microstoma* (Dujardin 1845). J Parasitol. 1963;49:305–7.

16. Hoff KJ, Lange S, Lomsadze A, Borodovsky M, Stanke M. BRAKER1: Unsupervised RNA-seq-based genome Annotation with GeneMark-ET and AUGUSTUS. Bioinformatics. 2016;32:767–9.

17. Bray NL, Pimentel H, Melsted P, Pachter L. Near-optimal probabilistic RNA-seq quantification. Nat. Biotechnol. 2016;34:525–7.

18. Emms DM, Kelly S. OrthoFinder: phylogenetic orthology inference for comparative genomics. Genome Biol. Genome Biology; 2019;20:1–14.

19. Waterhouse RM, Seppey M, Simao FA, Manni M, Ioannidis P, Klioutchnikov G, et al. BUSCO applications from quality assessments to gene prediction and phylogenomics. Mol Biol Evol. 2017;35:543–8.

20. Lynch M. The origins of genome architecture. Sinauer Associates Incorporated; 2007.

21. Canapa A, Barucca M, Biscotti MA, Forconi M, Olmo E. Transposons, genome size, and evolutionary insights in animals. Cytogenet Genome Res. 2016;147:217–39.

22. Volfovsky N, Haas BJ, Salzberg SL. Computational discovery of internal micro-exons. Genome Res. 2003;13:1216–21.

23. DeMarco R, Mathieson W, Manuel SJ, Dillon GP, Curwen RS, Ashton PD, et al. Protein variation in blood-dwelling schistosome worms generated by differential splicing of micro-exon gene transcripts. Genome Res. 2010;20:1112–21.

24. Brehm K, Frosch M, Jensen K. mRNA trans-splicing in the human parasitic cestode *Echinococcus multilocularis*. J Biol Chem. 2000;275:38311–8.

25. Boroni M, Sammeth M, Grossi Gava S, Andressa Nogueira Jorge N, Mara Macedo A, Machado CR, et al. Landscape of the spliced leader trans-splicing mechanism in *Schistosoma mansoni*. Sci Rep. Springer US; 2018;8:1–14.

26. Lee E, Harris N, Gibson M, Chetty R, Lewis SE. Apollo: a community resource for genome annotation editing. Bioinformatics. 2009;25:1836–7.

27. Stover NA, Katsanis N, Cavalcanti ARO. Spliced leader trans-splicing. Curr Biol. 2006;16:R8–9.

28. Krchňáková Z, Krajčovič J, Vesteg M. On the possibility of an early evolutionary origin for the spliced leader trans-splicing. J Mol Evol. Springer US; 2017;85:37–45.

29. Douris V, Telford MJ, Averof M. Evidence for multiple independent origins of trans-splicing in Metazoa. Mol Biol Evol. 2010;27:684–93.

30. Rossi A, Ross E, Jack A, Sánchez Alvarado A. Molecular cloning and characterization of SL3: A stem cell-specific SL RNA from the planarian *Schmidtea mediterranea*. Gene. 2014;533:156–67.

31. Koziol U, Jarero F, Olson PD, Brehm K. Comparative analysis of Wnt expression identifies a highly conserved developmental transition in flatworms. BMC Biol. BioMed Central; 2016;14:10.

32. Tandonnet S, Koutsovoulos GD, Adams S, Cloarec D, Parihar M, Blaxter M, et al. Chromosome-wide evolution and sex determination in the three-sexed nematode *Auanema rhodensís*. G3. 2019;:g3.0011.2019–20.

33. Rausch VR, Rausch RL. The karyotype of *Echinococcus multilocularis* (Cestoda: Taeniidae). Can. J. Genet. Cytol. 1981;23:151–4.

34. Pryde FE, Gorham HC, Louis E. Chromosome ends: all the same under their caps. Curr Opin Genet Dev. 1997;7:822–8.

35. Shelby RD, Vafa O, Sullivan KF. Assembly of CENP-A into centromeric chromatin requires a cooperative array of nucleosomal DNA contact sites. J. Cell Biol. Rockefeller University Press; 1997;136:501–13.

36. Hartley G, O’Neill R. Centromere repeats: hidden gems of the genome. Genes. 2019;10:223–22.

37. Melters DP, Bradnam KR, Young HA, Telis N, May MR, Ruby JG, et al. Comparative analysis of tandem repeats from hundreds of species reveals unique insights into centromere evolution. Genome Biol. BioMed Central Ltd; 2013;14:R10.

38. Bonfield JK, Whitwham A. Gap5--editing the billion fragment sequence assembly. Bioinformatics. 2010;26:1699–703.

39. Nielsen L, Edström JE. Complex telomere-associated repeat units in members of the genus chironomus evolve from sequences similar to simple telomeric repeats. Mol Cell Biol. American Society for Microbiology (ASM); 1993;13:1583–9.

40. Mason JM, Biessmann H. The unusual telomeres of *Drosophi/a*. Trends Genet. 1995;11:58–62.

41. McKinley KL, Cheeseman IM. The molecular basis for centromere identity and function. Sci Rep. Nature Publishing Group; 2015;17:16–29.

42. Saint-Leandre B, Levine MT. The telomere paradox: stable genome preservation with rapidly evolving proteins. Trends Genet. Elsevier Ltd; 2020;:1–11.

43. Olovnikov AM. A theory of marginotomy. The incomplete copying of template margin in enzymic synthesis of polynucleotides and biological significance of the phenomenon. J Theor Biol. 1973;41:181–90.

44. Victorelli S, Passos JF. Telomeres and cell senescence - size matters not. EBioMedicine. The Authors; 2017;21:14–20.

45. Tan TCJ, Rahman R, Jaber-Hijazi F, Felix DA, Chen C, Louis EJ, et al. Telomere maintenance and telomerase activity are differentially regulated in asexual and sexual worms. Proc Natl Acad Sci USA. 2012;109:4209–14.

46. Henikoff S, Ahmad K, Malik HS. The centromere paradox: stable inheritance with rapidly evolving DNA. Science. American Association for the Advancement of Science; 2001;293:1098–102.

47. Bracewell R, Chatla K, Nalley MJ, Bachtrog D. Dynamic turnover of centromeres drives karyotype evolution in Drosophila. eLife. 2019.

48. James K, Olson PD. The tapeworm interactome: inferring confidence scored protein-protein interactions from the proteome of *Hymenolepis microstoma*. BMC Genomics. 2020;:1–20.

49. Wences AH, Schatz MC. Metassembler: merging and optimizing de novo genome assemblies. Genome Biol. Genome Biology; 2015;16:1–10.

50. Koren S, Walenz BP, Berlin K, Miller JR, Bergman NH, Phillippy AM. Canu: scalable and accurate long-read assembly via adaptive k-mer weighting and repeat separation. Genome Res. 2017;27:722–36.

51. Chin C-S, Alexander DH, Marks P, Klammer AA, Drake J, Heiner C, et al. Nonhybrid, finished microbial genome assemblies from long-read SMRT sequencing data. Nat Meth. 2013;10:563–9.

52. Bankevich A, Nurk S, Antipov D, Gurevich AA, Dvorkin M, Kulikov AS, et al. SPAdes: a new genome assembly algorithm and Its applications to single-cell sequencing. J Comp Biol. 2012;19:455–77.

53. Kurtz S, Phillippy A, Delcher AL, Smoot M, Shumway M, Antonescu C, et al. Versatile and open software for comparing large genomes. Genome Biol. 2004;5:R12–9.

54. Staňková H, Hastie AR, Chan S, Vrána J, Tulpová Z, Kubaláková M, et al. BioNano genome mapping of individual chromosomes supports physical mapping and sequence assembly in complex plant genomes. Plant Biotechnol J. 2016;14:1523–31.

55. Dobin A, Davis CA, Schlesinger F, Drenkow J, Zaleski C, Jha S, et al. STAR: ultrafast universal RNA-seq aligner. Bioinformatics. 2013;29:15–21.

56. Flynn JM, Hubley R, Goubert C, Rosen J, Clark AG, Feschotte C, et al. RepeatModeler2: automated genomic discovery of transposable element families. 19:378. Available from: www.repeatmasker.org

57. Flynn JM, Hubley R, Goubert C, Rosen J, Clark AG, Feschotte C, et al. RepeatModeler2: automated genomic discovery of transposable element families. bioRxiv. Cold Spring Harbor Laboratory; 2019;19:856591.

58. Li L, Stoeckert CJ, Roos DS. OrthoMCL: identification of ortholog groups for eukaryotic genomes. Genome Res. Cold Spring Harbor Lab; 2003;13:2178–89.

59. Bernt M, Donath A, Jühling F, Externbrink F, Florentz C, Fritzsch G, et al. MITOS: Improved de novo metazoan mitochondrial genome annotation. Mol Phylogenet Evol. Elsevier Inc; 2013;69:313–9.

60. 14 GRCII42O211. BEDTools: a flexible suite of utilities for comparing genomic features. Bioinformatics. 2010;26:841–2.

61. McKenna A, Hanna M, Banks E, Sivachenko A, Cibulskis K, Kernytsky A, et al. The Genome Analysis Toolkit: a MapReduce framework for analyzing next-generation DNA sequencing data. Genome Res. Cold Spring Harbor Lab; 2010;20:1297–303.

62. Danecek P, Auton A, Abecasis G, Albers CA, Banks E, DePristo MA, et al. The variant call format and VCFtools. Bioinformatics. 2011;27:2156–8.

63. Orosová M, Špakulová M. Tapeworm chromosomes: their value in systematics with instructions for cytogenetic study. Folia Parasit. Folia Parasitologica; 2018;65:1–8.

64. Guo L, Accorsi A, He S, Guerrero-Hernández C, Sivagnanam S, McKinney S, et al. An adaptable chromosome preparation methodology for use in invertebrate research organisms. BMC Biol. BMC Biology; 2018;16:1–14.

65. Hopman AH, Ramaekers FCS, Speel EJ. Rapid synthesis of biotin-, digoxigenin-, trinitrophenyl-, and fluorochrome-labeled tyramides and their application for In situ hybridization using CARD amplification. J Histochem Cytochem. SAGE Publications; 1998;46:771–7.

66. Schindelin J, Arganda-Carreras I, Frise E, Kaynig V, Longair M, Pietzsch T, et al. Fiji: an open-source platform for biological-image analysis. Nat Meth. 2012;9:676–82.

67. Egger B, Bachmann L, Fromm B. Atp8 is in the ground pattern of flatworm mitochondrial genomes. BMC Genomics. BMC Genomics; 2017;18:1–10.

