## Supplementary material for "Complete representation of a tapeworm genome reveals chromosomes capped by centromeres, necessitating a dual role in segregation and protection": Add. File 2: Robot method

```
********** method-listing: Telomere TSA FISH 040419.IPM ***********

********** path: C:\INTAVIS\Software\InsituPro VSi 4.2\Methods\Methods
Slides\
** 08.04.2019 - 15:03:17
** Software : InsituProVSi
** Version : 29.05.2017
********** configuration: InsituPro VSi-TSd.IPC **********
********** path: C:\INTAVIS\Software\InsituPro VSi 4.2\
* XYZRoboter - Gilson 223

- Drain / Rinse / Home  X / Y / Z -
Home  :                      X:270 Y:205 Z:0    F:0      0.1mm
Drain :                      X:270 Y:205 Z:300  F:300    0.1mm
Rinse :                      X:275 Y:530 Z:1080 F:1080   0.1mm
- change Needle -
position :                   X:0   Y:0   Z:0    F:0      0.1mm
* Dilutor - Gilson 402
- Identification : -
Alias : [empty => unit-nb]

- Configuration -
dilutor syringe:              10000
transfertube:                 10100                     ul

- Pipetting -
reservoir aspiration :        30                        ml/min
MANUAL prime speed:           72                        ml/min
Reservoir-Volume:             2000                      ml
aspiration speed LOW:         10                        ml/min
aspiration speed MEDIUM:      15                        ml/min
aspiration speed HIGH:        20                        ml/min
dispense speed LOW:           10                        ml/min
dispense speed MEDIUM:        15                        ml/min
dispense speed HIGH:          20                        ml/min
inside rinse vol. NORMAL:     700                       µl
inside rinse vol. INTENSIVE:  1400                      µl
outside rinse vol. NORMAL:    700                       µl
outside rinse vol. INTENSIVE: 1400                      µl
rinse dispense speed LOW:     20                        ml/min
rinse dispense speed HIGH:    30                        ml/min

(System: Slides)
* ThermContr - WEST 6100+
- Identification : -
Alias : [empty => unit-nb]
- Method -
- Contacts -
output cool-fan :             4                         1..4

********** method *****************************
1     Module     Starting Method
1.1   XYZCheck
1.2   SetTempReg  to : OFF
1.3   PrimeNeedle 12000µl
1.4   PrimePort   10000µl Port 1->Drain
1.5   PrimePort   10000µl Port 2->Drain
1.6   PrimeTub    60000µl
```

```
...
2     Module      Hybridization
2.1   SetTempReg  to : 70°C
2.2   IncubateTS  00:10 250µl Hyb Mix->Slides
2.3   SetTempReg  "to : 70°C, <30min"
2.4   IncubateTS  00:10 250µl Hyb Mix->Slides
2.5   PrimeTub    60000µl
2.6   IncubateTS  00:30 200µl Probe->Slides
2.7   SetTempReg  to : RT
2.8   Wait  12 h
2.9   PrimeTub    60000µl
2.10  IncubateTS  00:05 250µl 2x SSC->Slides 6x
2.11  SetTempReg  to : OFF
2.12  PrimeTub    60000µl
2.13  IncubateTS  00:05 250µl 0.5x SSC->Slides 6x
2.14  IncubateTS  00:05 250µl TNT->Slides 6x
2.15  PrimeTub    60000µl
...
3     Module      AB incubation
3.1   IncubateTS  00:15 250µl Block->Slides 2x
3.2   IncubateTS  02:00 250µl AB1(aDIG-POD->Slides
3.3   IncubateTS  00:05 250µl TNT->Slides 6x
3.4   IncubateTS  00:05 250µl PBST->Slides 2x
3.5   IncubateTS  00:05 250µl PBS Imid->Slides 2x
3.6   IncubateTS  00:05 250µl TSA1->Slides
3.7   IncubateTS  00:05 250µl PBST->Slides 6x
3.8   IncubateTS  00:05 250µl TNT->Slides 6x
3.9   IncubateTS  00:15 250µl DAPI->Slides 2x
3.10  IncubateTS  00:05 250µl TNT->Slides 2x
3.11  PrimeTub    60000µl
...
4     Module      Finishing method
4.1   PrimeTub    60000µl
4.2   RinsePort   10000µl Reservoir->Port 1
4.3   RinsePort   10000µl Reservoir->Port 2
4.4   PrimeNeedle 12000µl
4.5   SetTempReg  to : OFF
...
...
********** end of method *** ******************
```
