## Supplementary material for "Complete representation of a tapeworm genome reveals chromosomes capped by centromeres, necessitating a dual role in segregation and protection": Add. File 3: Supp. Figures S1-10

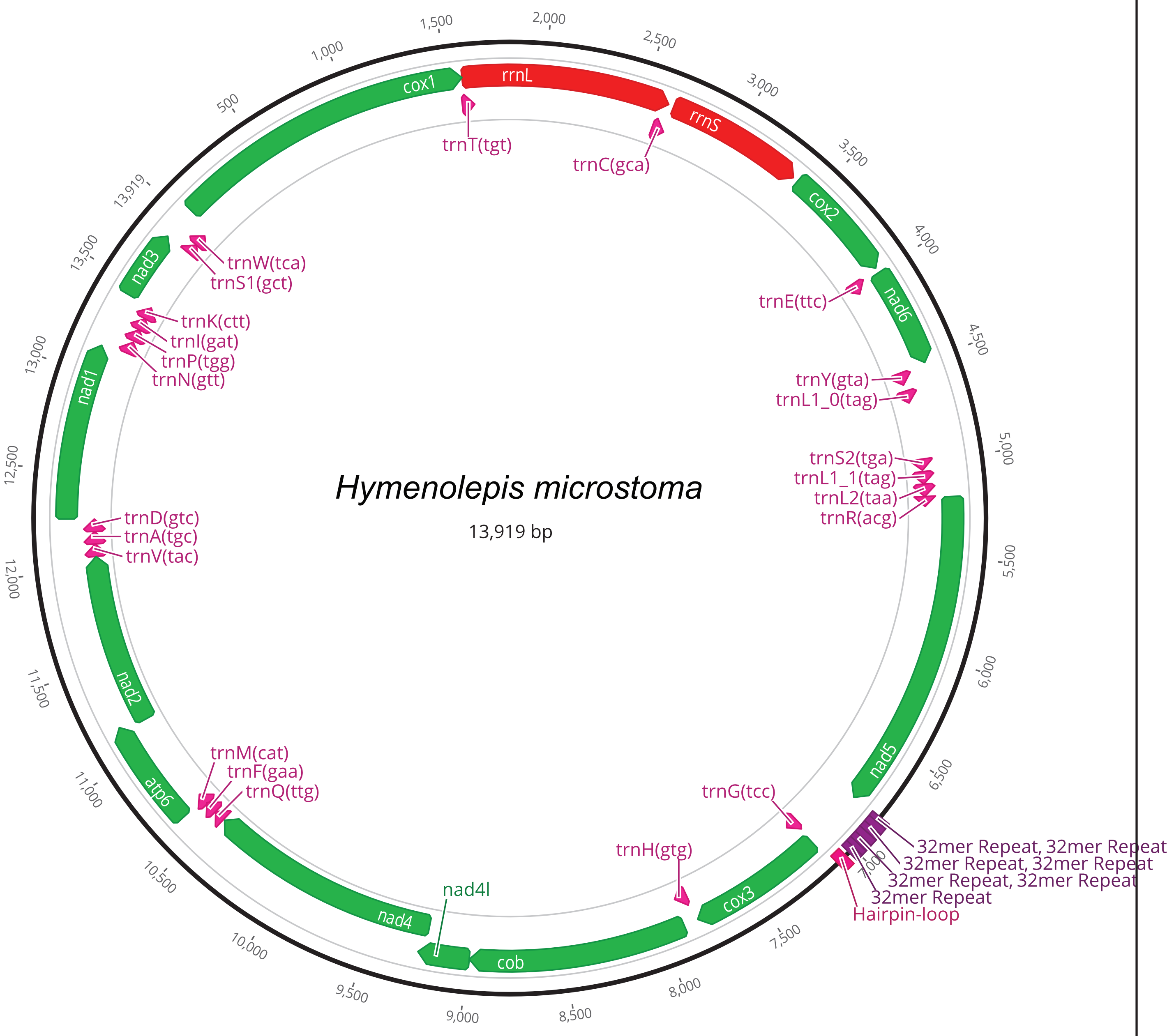


**Supplementary Fig. S1.** Mitochondrial genome. The 13,919 bp *Hymenolepis microstoma* mitochondrial genome was re-assembled from both short and long-read data, yielding over 1000x coverage. The new assembly resolved the full length of a region involving a tandemly repeated 32 bp motif (cf. GenBank accession AP017665.1). This region is identified as one of three origins of replication-heavy strand (OH-a) by MITOS [59] and an adjacent hairpin-loop region as the origin of replication-light strand (OL). Gene order of ribosomal and protein-coding genes is consistent with the hypothesized ground-plan for the mitogenomes of parasitic flatworms as is the absence of the atp8 gene [67].

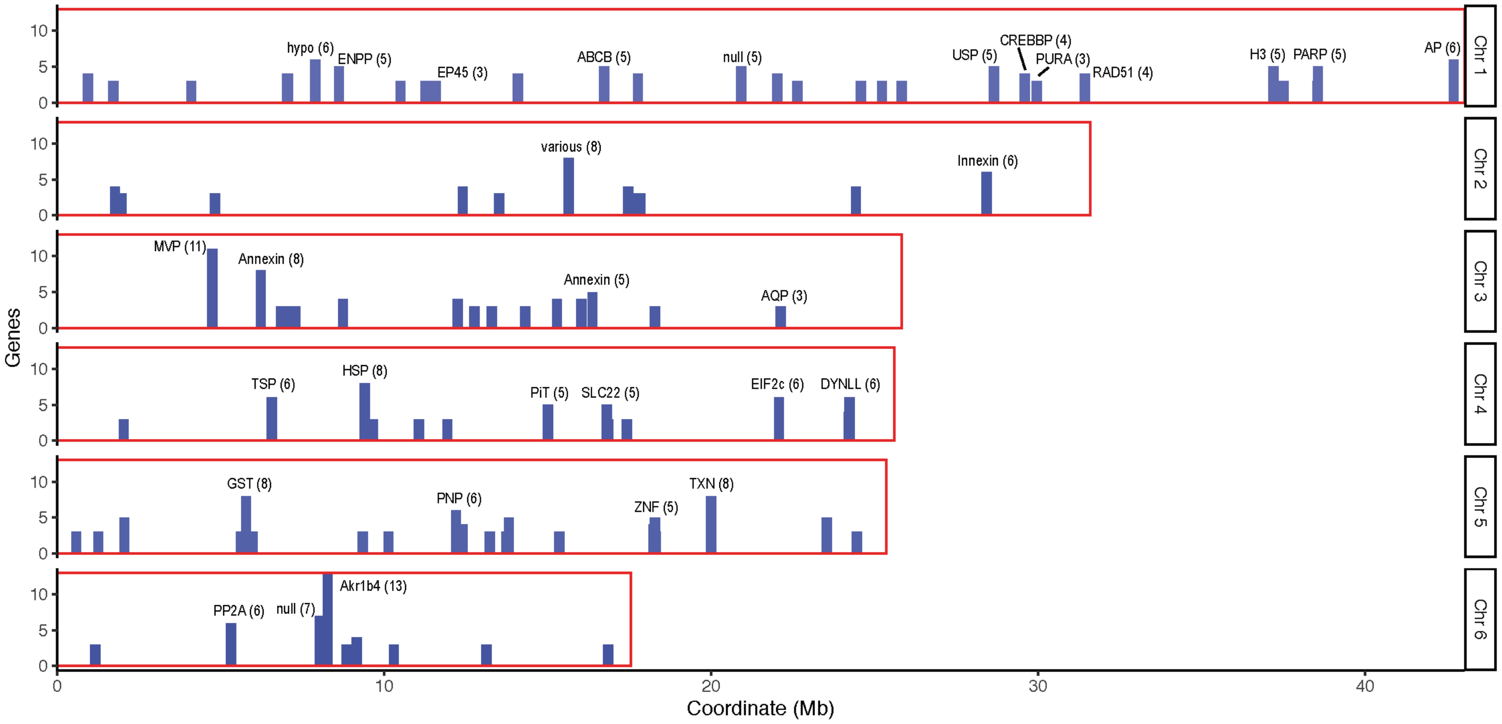


**Supplementary Fig. S2.** Repeat hotspots. Chromosomal positions of paralogous gene arrays. Abbreviations: ABCB: ATP binding cassette subfamily B; Akr1b4: Aldo keto reductase family 1 member B4; AP: Alkaline phosphatase; AQP: Aquaporin 4; CREBBP: CREB binding protein; DYNLL: Dynein light chain; EiF2c: Eukaryotic translation initiation factor 2c; ENPP: Ectonucleotide pyrophosphatase:phosphodiesterase; EP45: Estrogen regulated protein EP45; GST: Glutathione S transferase; H3: Histone H3; HSP: heat shock protein; hypo: hypothetical protein; MVP: Major vault protein; PARP: Poly [ADP-ribose] polymerase; PiT: Phosphate transporter; PNP: Purine nucleoside phosphorylase; PP2A: Serine:threonine protein phosphatase 2A; PURA: PUR alpha protein; USP: Universal stress protein; RAD51: DNA repair protein RAD51 homolog; SLC22: Solute carrier family 33; TSP: Tetraspanin; TXN: Thioredoxin; ZNF: zinc finger protein.

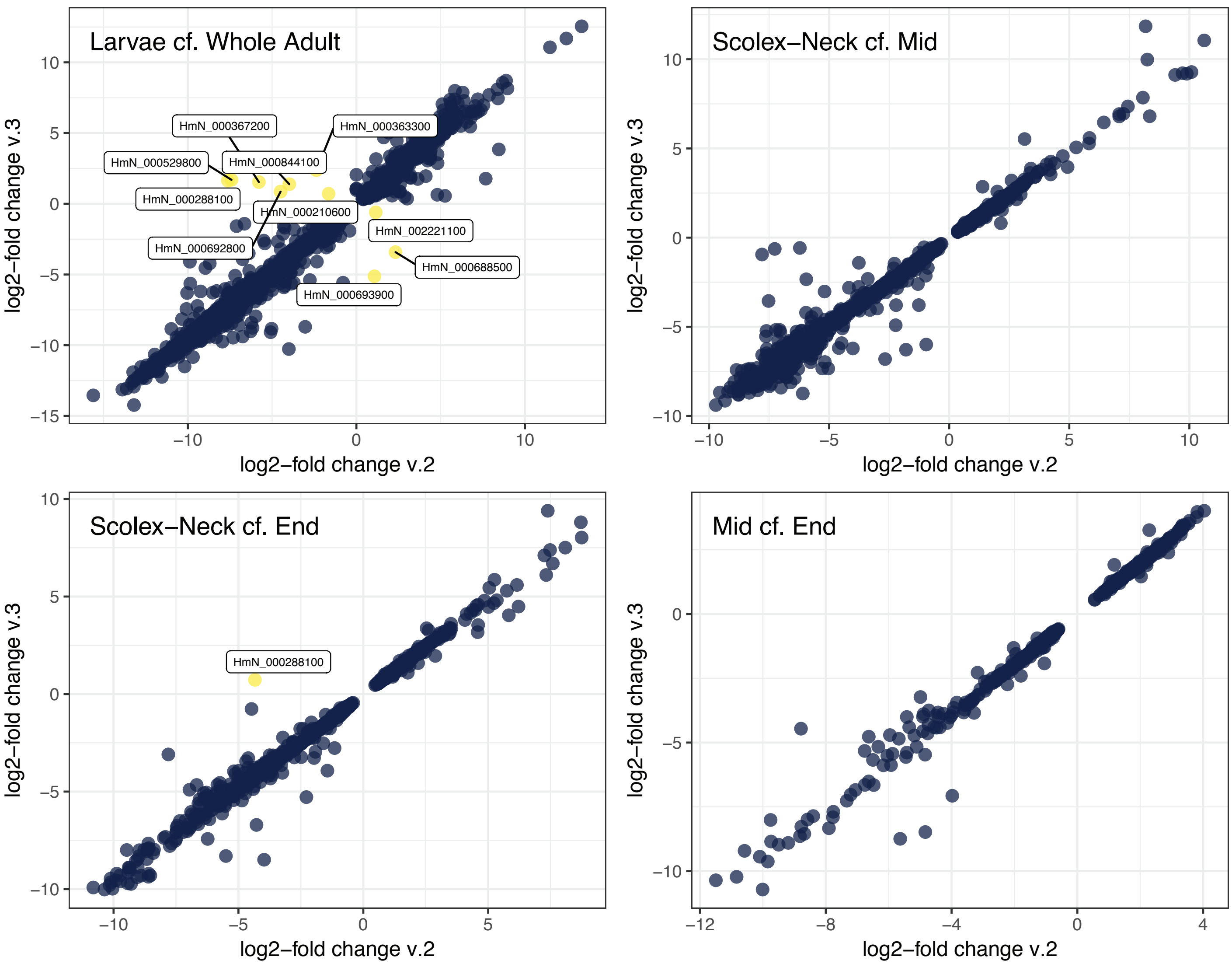


**Supplementary Fig. S3.** Comparison of differentially expressed genes estimated from RNA-seq counts aligned to the v2 and v3 assemblies. Plots of log2-fold change show highly linear relationships across all sample comparisons, corroborating previous findings [8]. Only 11 genes (yellow), all with small fold-change values, were found to reverse directionality between assembly versions.

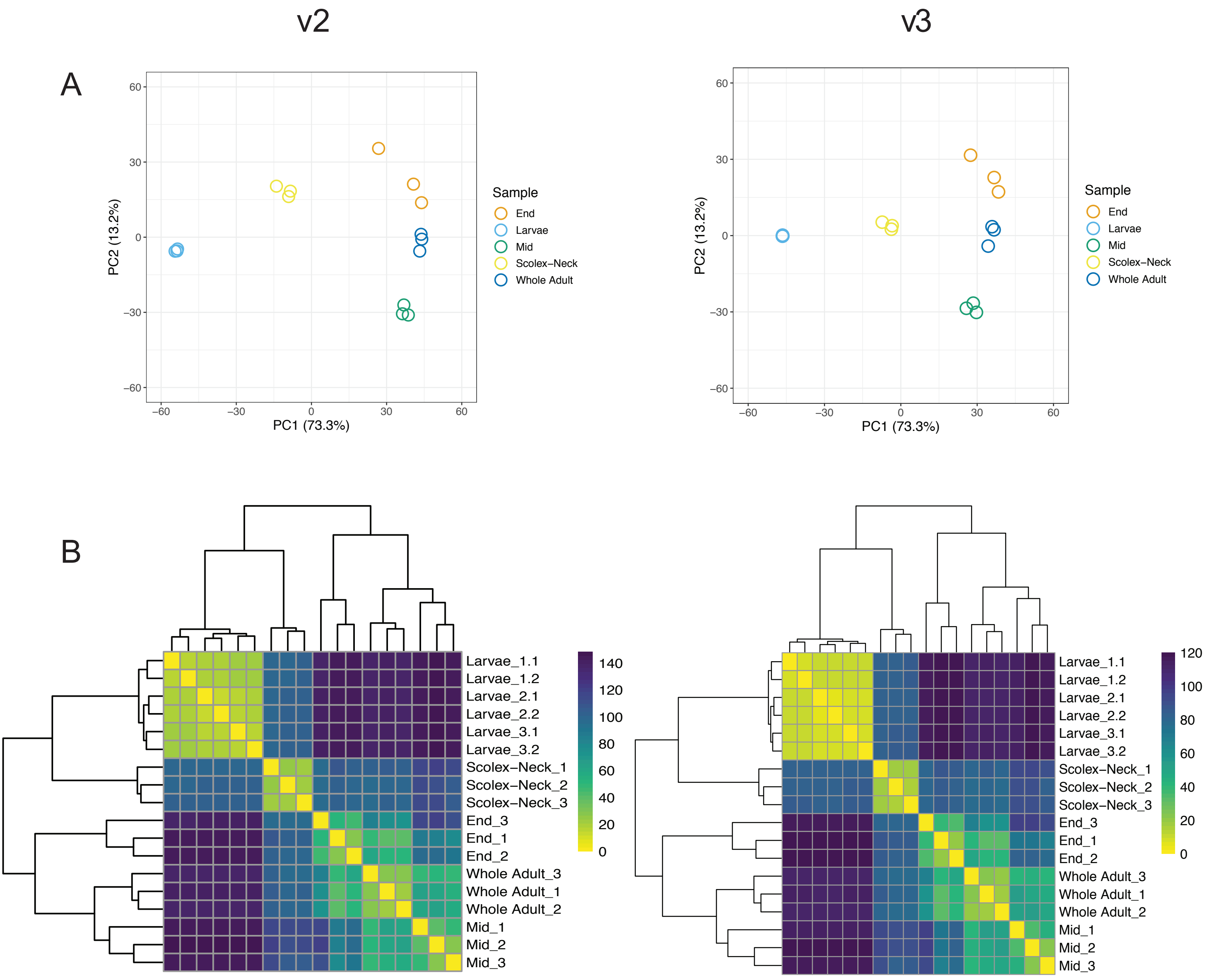


**Supplementary Fig. S4.** Comparison of RNA-seq sample counts against the v2 and v3 assemblies and gene models. Principle component analyses (**A**) show tight clustering of sample replicates based on counts using both assemblies, while in the v3 (right) the Larvae, Scolex-Neck and Whole Adult samples are arrayed only along PC1, with the transcriptome of the Scolex-Neck mid-way between those of the Larvae and Whole Adult samples. The Mid and End samples are further differentiated from the other samples along PC2. Heatmap clustering (**B**) shows that the transcriptome of the Scolex-Neck region is more similar to that of mid-metamorphose larvae than to middle or end regions of the adult worm, as discussed in [8].

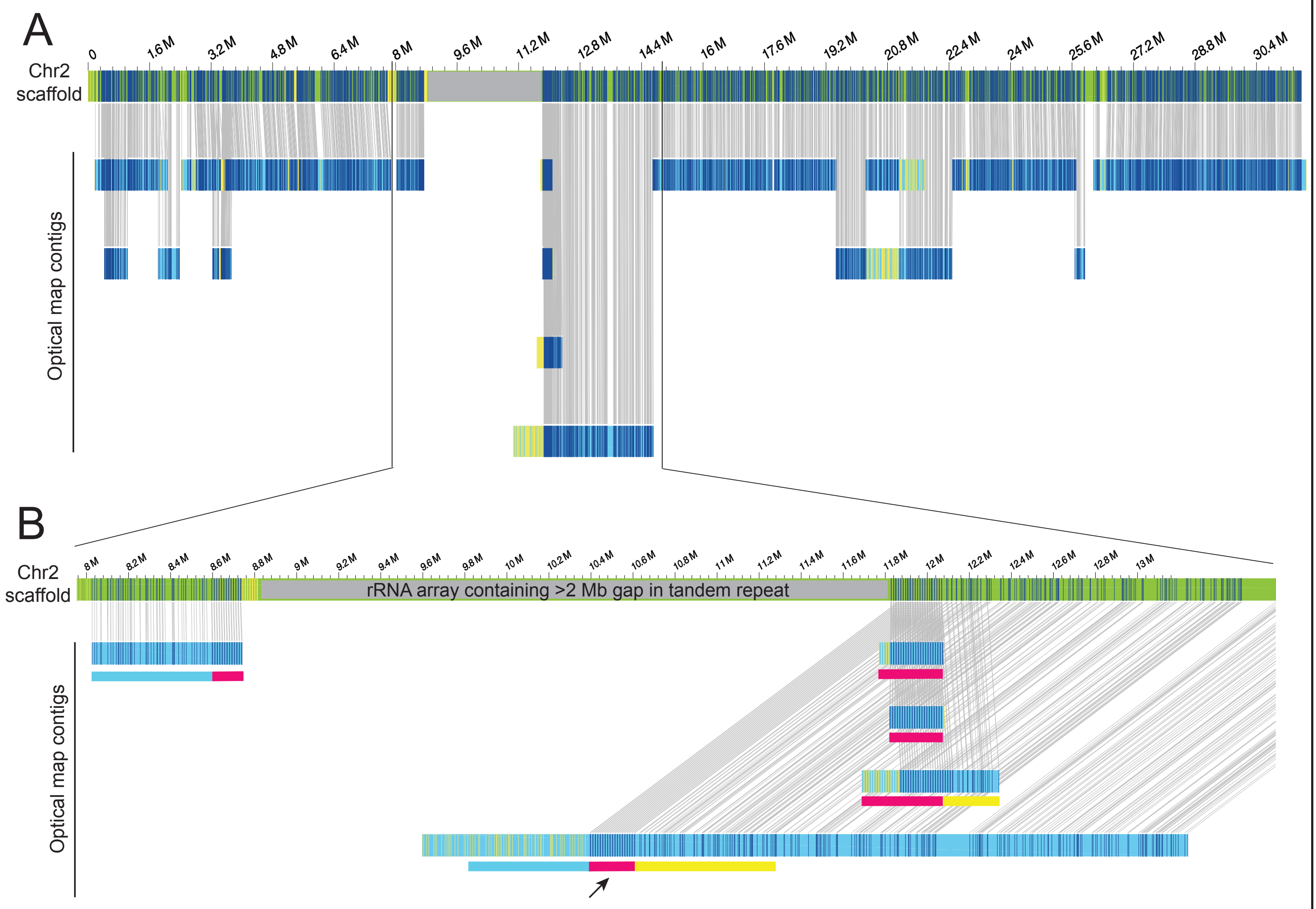


**Supplementary Fig. S5.** Optical map contigs aligned to the genome assembly of the rRNA repeat array. Five contigs from the optical map are shown with the segment that aligns to the sequenced repeat indicated by coloured bars. The largest map contig (arrow) represents one haplotype containing the rRNA tandem repeat (pink bar) as well as the left (blue bar) and right (yellow bar) flanking regions. Other optical map contigs either contain the repeat together with either 5’ or 3’ flanking region, and likely represent an alternative haplotype, or have an insufficient amount of unique sequence to unambiguously determine their position within the repeat array.

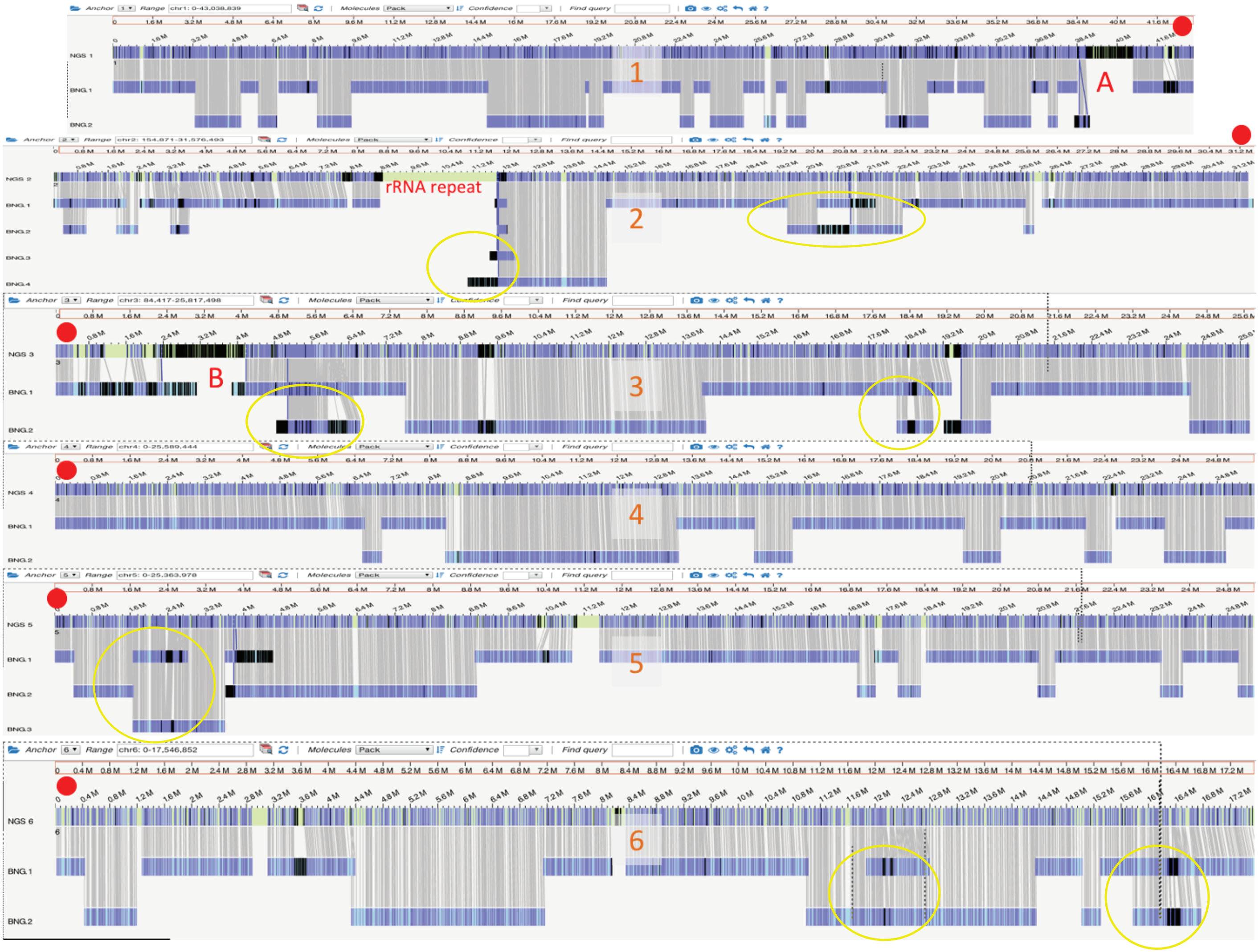


**Supplementary Fig. S6.** Whole genome optical maps aligned to v3 assembly. Circled regions show where optical map data indicate alternative haplotypic versions. Regions labelled A and B, together with the rRNA array, represent the largest repeat regions where haplotype differences could account for visible length differences in sister chromatids (see text). Chromosomes are numbered and the positions of the telomeric repeats indicated by red dots.

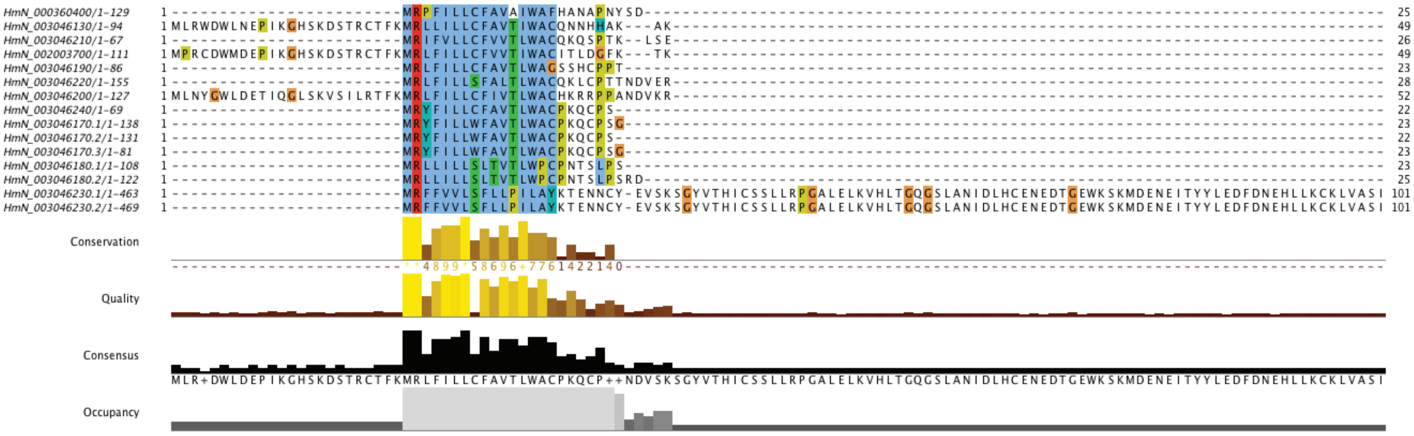


**Supplementary Fig. S7.** Alignment of the N-terminal regions encoded by a tandem array of micro-exons genes located on chromosome 6. The shared amino acid motif (consensus MRLFILLCFAVTLWAC) indicates that this gene array evolved through tandem duplication.

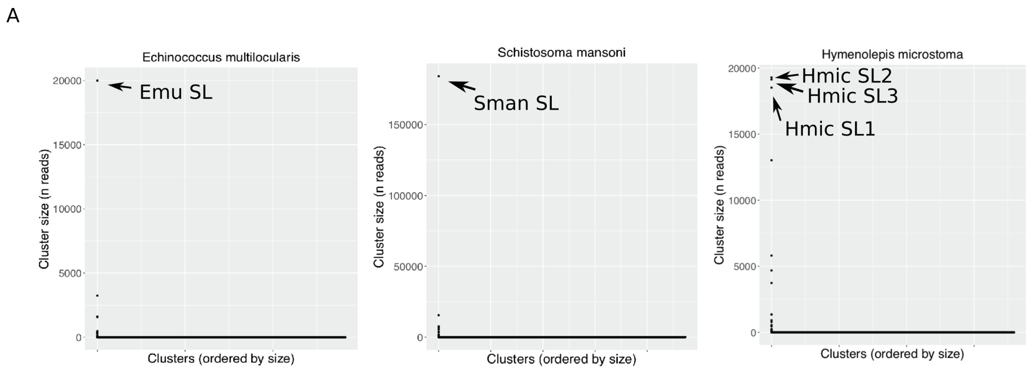


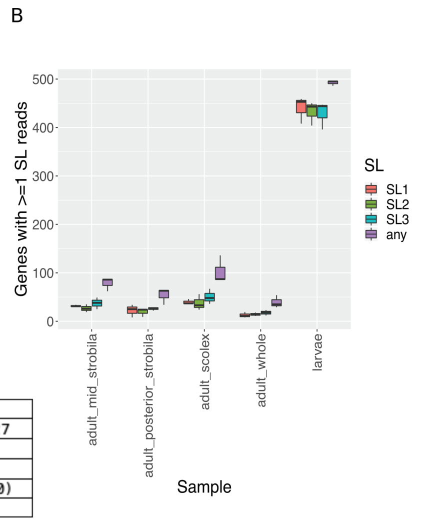


| SL | Sequence (derived from clipped reads) | Genomic location |
|---|---|---|
| Hmic SL1 | CCGGTCTTACCATAAAACTTGTAT | HMN_01_pilon:9215213..9215235 , HMN_01_pilon:9216005..9216027 |
| Hmic SL2 | CCGGTCTTACCTTGCAATTTTTGTAT | HMN_03_pilon:20837621..20837646 |
| Hmic SL3 | AATCGGTCTTACTGTACTAACTTGTAT | HMN_01_pilon:5649153..5649179 |
| Sman SL | CACGGTTTTACTCTTGTGATTTGTTGCAT | 118 loci on 5 scaffolds (of which 108 within SM_V7_6:6847000..6942300) |
| Emu SL | TCGGTCCTTACCTTGCAGTTTTGTAT | 67 loci on 8 contigs/scaffolds |

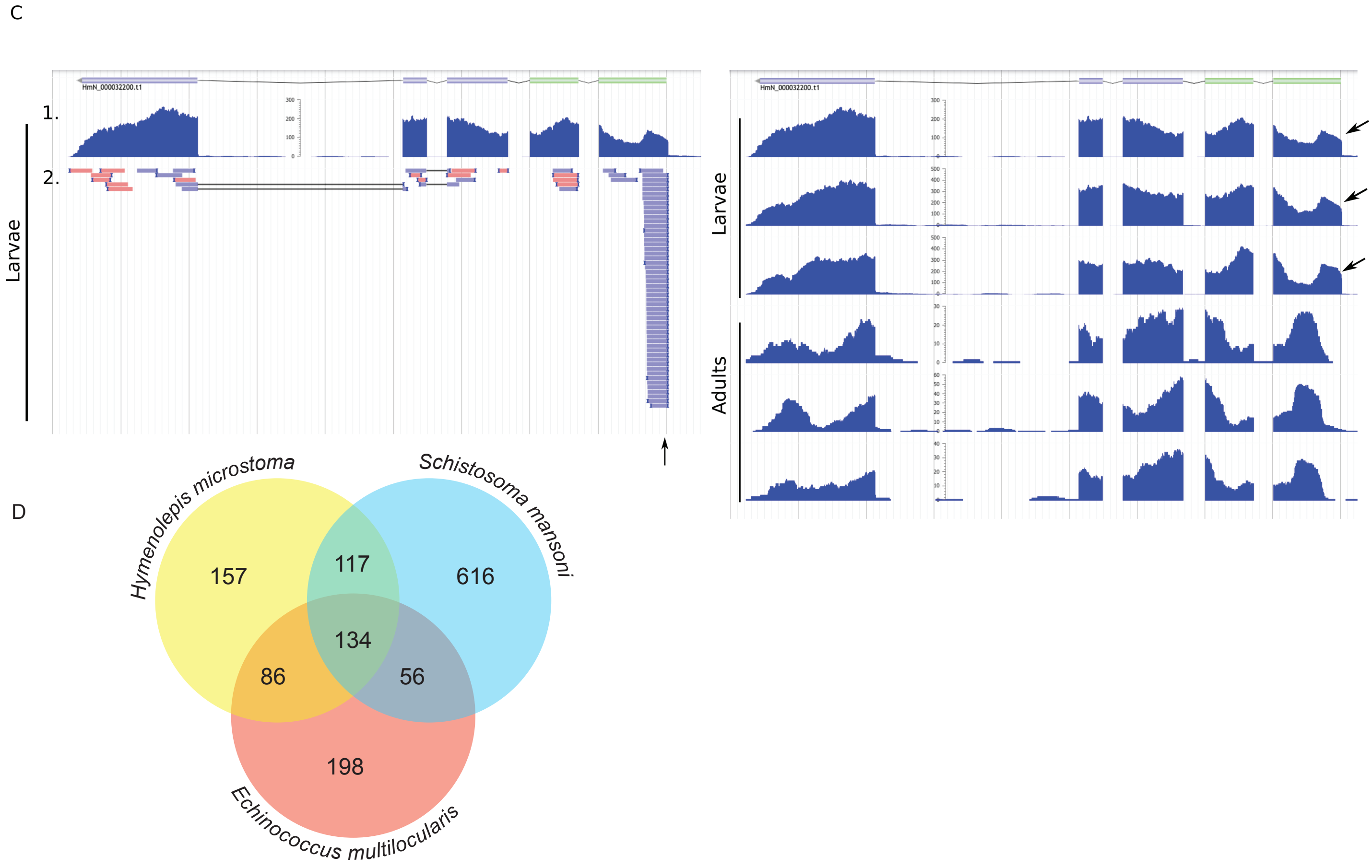


**Supplementary Fig. S8.** Spliced leader trans-splicing. (**A**) Clustering of sequences soft clipped from aligned RNASeq reads. The most abundant clusters represent known (*E. multilocularis*, *S. mansoni*) or candidate (*H. microstoma*) splice leader (SL) sequences which are given in the table below. (**B**) Prevalence of trans-splicing in different life stages and regions of the adult worm. Genes were considered trans-spliced if > 10 SL reads (SL1, SL2 or SL3) aligned across all libraries analysed. Of these genes, plot represents instances of at least one SL read aligning in each sample. Note that there almmost 5x as many genes trans-spliced in larvae than in the adult samples. Three replicates per sample. (**C**) An example of a gene (HmN_000032200) that is trans-spliced in larval but not adult samples, visualised using Apollo. Left: track 1 shows a coverage plot of all aligned reads; track 2 represents alignments of uniquely-mapping soft-clipped reads (soft clipping represented by a thick blue bar at the end of the read). Arrow indicates accumulation of soft clipped reads at proposed SL-acceptor site. Right: Coverage plots of all aligned reads in three larval and three adult libraries. Arrows indicate proposed SL-acceptor sites present in the larval but not adult libraries. (**D**) Venn diagram of trans-spliced orthogroups shared between parasitic flatworms.

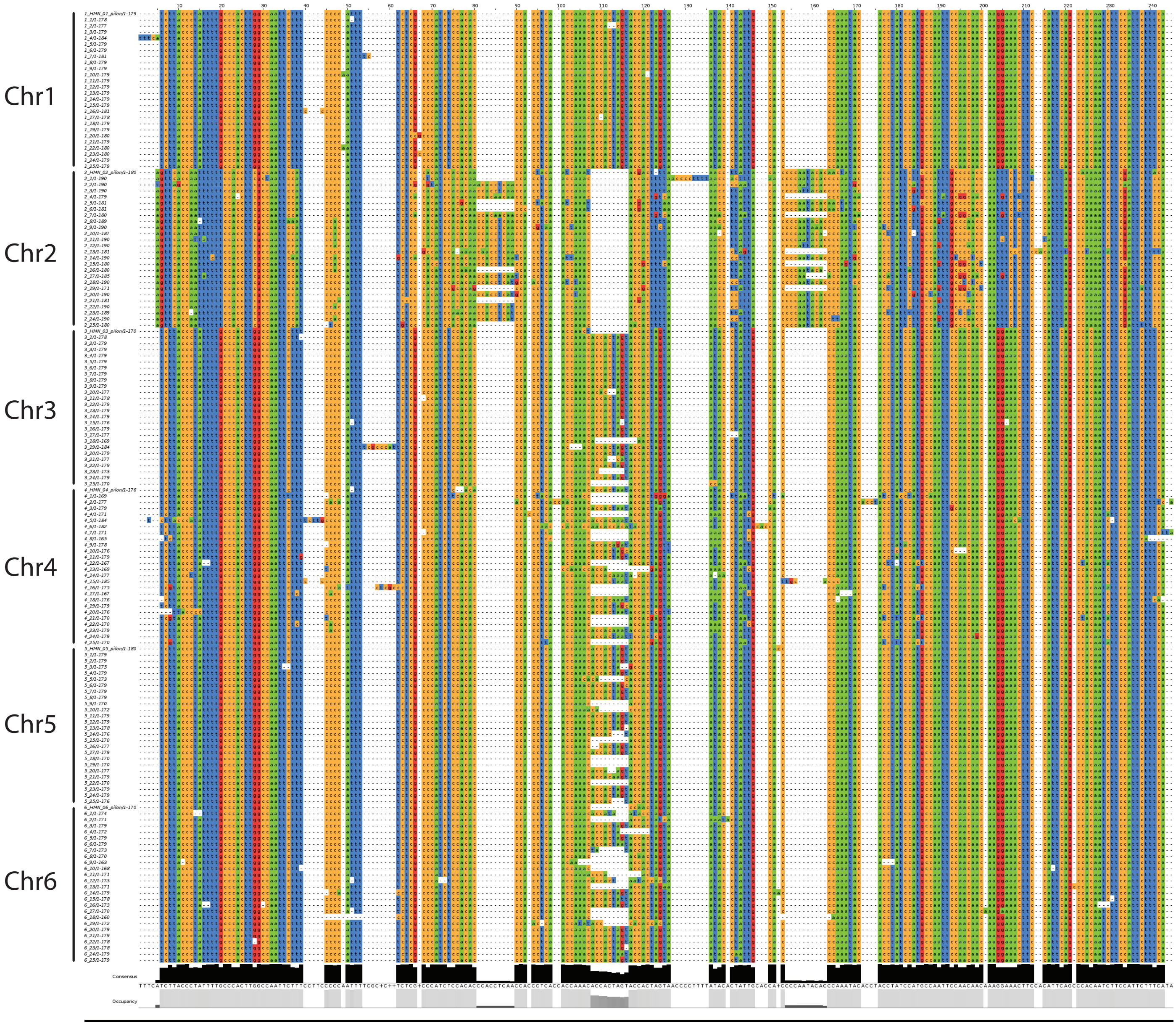


**Supplementary Fig. S9.** Multiple alignment of the terminal centromeric repeats of each chromosome. 26 consecutive repeat copies were taken from a single location at the end of each of the six chromosomes in turn and aligned in order (top 26 = Chr1, next 26 = Chr2 etc.). Strong conservation of the 179mer centromeric repeat is seen across all chromosomes except Chr2 which shows a second novel repeat type. However, searching within the whole of the Chr2 repeat array shows that the 'canonical' 179mer observed in the other aligned reads is found with 100% coverage and identity. The terminal array on Chr2 is also much larger than those of the other chromosomes and is interspersed with various other repeats not shown here. Full assembly of the Chr2 terminal array is not resolvable without longer sequencing reads. Notably, when the centrosomal repeat arrays are oriented at the same end of each chromosome their sequences are found to be in alignment.

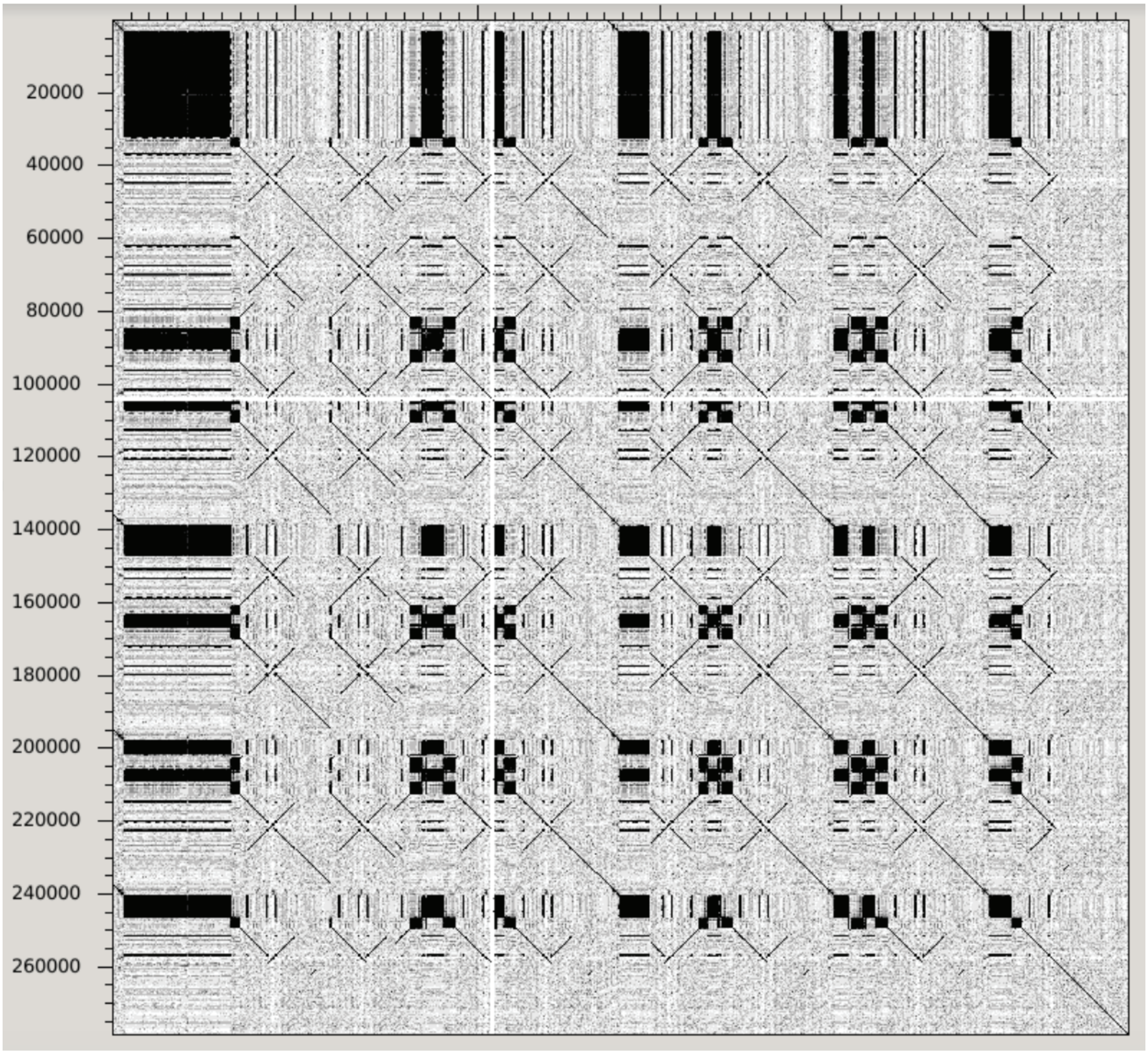


**Supplementary Fig. S10.** The terminal centromeric repeat of chromosome 2. A dotter plot shows that the centromeric repeat not only contains a second dominant repeat motif but is also interspersed with other repetitive elements, unlike the other chromosomes that exhibit a tandem array comprised entirely of the novel 179mer. Within the interstitial sequences we find the top blastx hit to Gag-Pol polyprotein, indicating the centromere has been invaded by transposable elements.
